# Ancient metagenomics reveals subglacial microbiomes driven by oxygen availability

**DOI:** 10.64898/2025.12.03.692186

**Authors:** Bianca De Sanctis, Nicholas B. Dragone, Ciara Wanket, Clifton P. Bueno de Mesquita, Sarah Crump, Gavin Piccione, Berkhashni Nirula, Halle Bender, Russell Corbett-Detig, E. Troy Rasbury, Graham Edwards, Abigale Hawthorn, John J. Welch, Alexandra Rouillard, Beth Shapiro, Jill Mikucki, Terrence Blackburn

## Abstract

Beneath Earth’s glaciers and ice sheets lies an aquatic realm where ice, water, rock, and microbial life interact, driving chemical reactions that can collectively influence the global carbon cycle, polar oceans, and climate. Efforts to describe subglacial microbiomes have been limited by the challenge of cleanly drilling through hundreds of meters of ice, such that only a few sites have ever been directly sampled. Here we use ancient metagenomics to present the first spatiotemporal characterization of subglacial bacteria and archaea. We extracted DNA from 25 subglacial precipitate samples, sedimentary accumulations of minerals that form in subglacial waters prior to exposure on the surface. The precipitates studied here formed between 16,000 and 570,000 years ago beneath the Antarctic and Laurentide Ice Sheets. We show that postmortem DNA damage patterns can reliably distinguish between ancient subglacial and modern surface taxa, and that this approach can enable reconstruction of subglacial microbiomes across poles and ice ages. Our analysis suggests that subglacial microbiomes are dominated by chemolithoautotrophs, ultra-small microbes, and taxa closely related to those found in deep subsurface or extreme cold and hypersaline environments. These microbiomes split into two distinct clusters distinguished by oxygen availability and redox conditions, irrespective of geography or age. Geochemical measurements of subglacial redox state, measured either indirectly via precipitate calcite Fe and Mn concentrations or directly via water reduction potential, reproduce these same two clusters exactly. Our findings describe how subglacial water redox states are held in balance by microbes, hydrology, and oxygen input from fresh subglacial meltwater, that we interpret to be controlled by the ice sheet response to past climate variations.

## 1 Main

Glaciers cover roughly 10% of Earth’s terrestrial surface, but the aquatic environments hidden underneath them are some of the most inaccessible on the planet. These subglacial ecosystems harbor polyextremophilic microbes whose metabolisms drive biogeochemical cycles that influence greenhouse gas emissions, enhance bedrock weathering, and promote nutrient export to polar oceans [1, 2]. Ice sheets respond to climate forcing on centennial to orbital timescales, producing variations in subglacial water chemistry that impact the oxygen availability and energy sources for microbes, constraining their structure and function [1, 3, 4]. Understanding the complex interactions between subglacial microbiomes, biogeochemical processes, and ice sheet hydrology is therefore critical for predicting the role of these systems in our changing world. Beyond Earth, subglacial environments serve as analogs for potential subsurface ocean habitats on icy moons such as Europa and Enceladus [5].

Despite their importance, clean subglacial samples remain exceedingly rare. Accessing subglacial lakes requires considerable effort and logistical resources such as transporting large drilling equipment and fuel, establishing large field camps in remote regions, and microbiological studies further demand clean access approaches where drilling equipment must be decontaminated [6, 7]. In Antarctica, which harbors at least 600 subglacial lakes and by far the most subglacial water on Earth [1], clean samples have only been collected from 3 sites: the neighbouring West Antarctic Subglacial Lake Whillans (SLW) and Subglacial Lake Mercer (SLM) [8, 9, 10, 11], and from an outlet glacier of the East Antarctic ice sheet, an aquifer below Taylor Glacier, referred to as Blood Falls Brine (BFB) [12, 13]. Both SLW and SLM are oxygen-rich, hydrologically active systems supporting aerobic phylotypes, whereas BFB is suboxic, with more metabolically plastic phylotypes [8, 9, 10, 11, 12, 13]. Given these differences, it remains challenging to define the typical structure and function of subglacial microbiomes, creating significant uncertainty for models of continental-scale nutrient fluxes and climate-sensitive gases.

Subglacial precipitates are sedimentary accumulations of minerals that can act as time capsules of past subglacial waters and their contents (see Figure 1A) [14, 15]. These precipitates form as a consequence of bedrock dissolution by subglacial waters followed by salt concentration to the point of mineral–calcite or opal-saturation. Precipitate formation can be accurately dated (via ^234^U −^230^ Th; see Methods) and combined with polar climate records to study how climate and ice sheet dynamics influence subglacial waters. For example, cycles in subglacial water chemistry recorded by precipitate mineralogy and isotopic results have been linked to the melt-freeze cycles of ice sheets, providing insights into the ice sheet response to climate change [15]. Subglacial precipitates can be exposed to the surface when: (1) ice retreats and uncovers underlying deposits, for example during recent interglacial warming; or (2) they are frozen into basal ice which is pushed vertically upwards by bedrock protrusions and lost via sublimation in blue ice regions (see Figure 1B) [16, 17]. Previous work constrained precipitate formation to within ∼10 km of the exhumed moraines using ice velocity and radioisotopic ages [15]. Calcite and opaline silica, of which these precipitates are composed, are two minerals known to be highly conducive to ancient DNA preservation [18, 19]. Key drivers of DNA degradation are oxidative, thermal, and hydrolytic stressors, and UV radiation [20, 21, 22, 23], all of which may be limited in these dark, cold subglacial environments, especially if water penetration is limited after precipitate formation.

**Figure 1:**
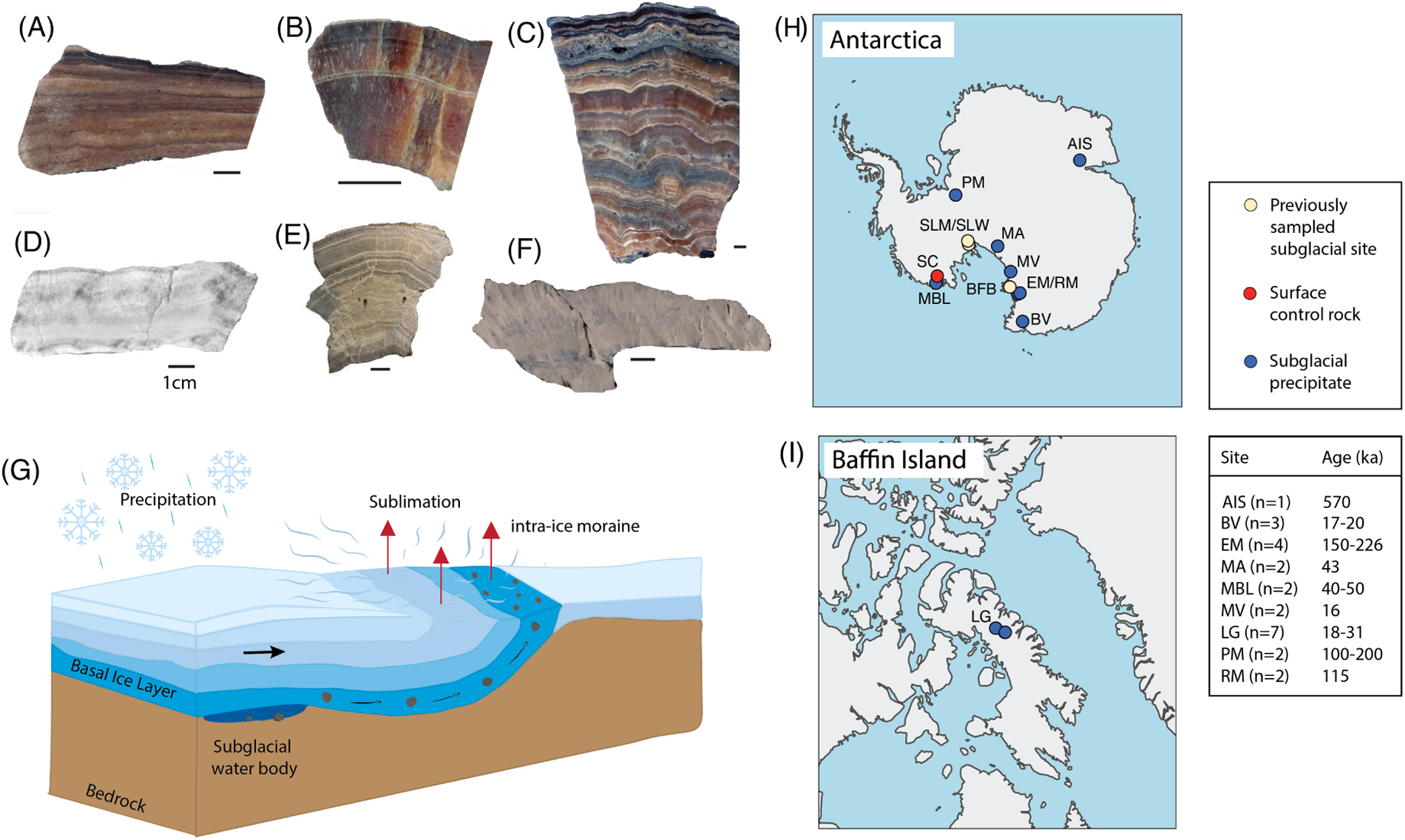
Formation and origin of subglacial precipitates. (A-F) Representative rock precipitates from Antarctica. The structure, color and composition of samples varies with each site. Samples LG1(A), BV1 (B), MA1 (C) are representative of this studies “reduced” cluster. Note the orange to red calcite color. Samples AIS1 (D), RM1 (E), MBL1 (F) are representative of this studies “oxidized” cluster. Scale bars are 1cm. (B) Cartoon schematic of precipitate formation (in subglacial water body), freeze-in addition to a moving basal ice layer, before exhumation at supgraglacial moraines. Black arrow notes direction of ice motion (C) and (D) show sample locations on Antarctica and Baffin island, respectively. Site codes are: LG = Laurentide Gneissic bedrock, AIS = Amery ice shelf, PM = Pensacola Mountains, BV = Boggs Valley, EM = Elephant Moraine, RM = Reckling Moraine, MV = Magnus Valley, MBL = Marie Byrd Land, MA = Mount Archernar, and SC = Surface control, which is from Vivan Nunatak (also in Marie Byrd Land). Previous cleanly sampled subglacial sites are SLM = Subglacial Lake Mercer, SLW = Subglacial Lake Whillans, BFB = Blood Falls Brine. Ages and sample counts are shown for the subglacial precipitate sites.

Here, we shotgun sequenced ancient DNA trapped inside subglacial precipitates originating from below the Antarctic and Laurentide ice sheets. Despite the absence of the Laurentide ice sheet today, preserved subglacial features reveal a multitude of past subglacial lakes suggesting this ice sheet, like Earth’s current ice sheets, harbored an active subglacial hydrologic system in the past [24]. Photographs of six subglacial precipitates used in this study are shown in Figure 1A, and sample locations are shown in Figure 1H-I. Our sample set includes 17 previously dated precipitate samples [14, 15, 25, 26] and 1 newly dated (reported here, Supplementary Data 1.4), that span the continent of Antarctica and range in formation age from 20,000-570,000 years old, 7 precipitates from two locations near the Barnes Ice Cap on Baffin Island dated between 18,000-31,000 years old [24], and a calcite sample that formed on an Antarctic rock surface used as a control (Figure 1). Using postmortem DNA damage to distinguish ancient subglacial from modern surface taxa, we identify two subglacial microbial clusters, a partitioning that we show is driven by oxygen availability. Within these clusters, we find that the subglacial microbiomes are for the most part consistent across both poles and at least the last 570,000 years, even down to the genus level. The same clusters can be reproduced independently via geochemistry, either by precipitate manganese and iron concentrations or by water reduction potential. These results closely link geochemistry and microbiome composition in subglacial ecosystems, providing an evidence-based framework for subglacial conditions globally and potentially on other planets.

## 2 Postmortem damage effectively separates subglacial from surface taxa

We shotgun sequenced 25 ancient subglacial precipitate samples and a modern Antarctic surface control calcite (SC1). Reads were aligned against all representative genomes in the GTDB v226.0 archaea and bacteria genome catalogue [27], decontaminated against the negative controls (extraction and library blanks) and common eukaryotic contaminants, and assigned to the taxonomic node representing the lowest common ancestor of their alignments (see Methods).

Each of the precipitates has potentially been exposed to the surface microbiome for millennia, and it would be unreasonable to assume that all of the microbial DNA we sequenced originated from subglacial waters (as was done in [28]; see Supplementary Section 5.2). To separate subglacial from surface taxa, we use postmortem damage, a hallmark signature of ancient DNA, which is characterized by an increased frequency of deamination or C-to-T misincorporations on the 5’ and 3’ ends of single-stranded DNA fragments following cellular death [29]. To visualize the hierarchical taxonomic structure and associated damage patterns in each sample, we generated circular Krona plots (Figure 2) [30], with segments coloured by postmortem damage (5’ C-to-T frequency), and in which inner rings represent coarser taxonomy and outer rings show finer taxonomic resolution. Most samples show a mixture of minimally damaged surface taxa, and damaged ancient subglacial taxa (e.g. LG6, 24ky old, Figure 2B). The surface control rock (Figure 2A) and the negative controls yielded entirely undamaged DNA. Some precipitates, such as PM1 (Figure 2C), which is 200ky old, contain almost exclusively highly damaged, ancient DNA. Two precipitate samples, EM1 and PM2, contain the lowest number of assigned reads out of all the samples (Supplementary Figure 2), and little to no discernible ancient DNA.

**Figure 2:**
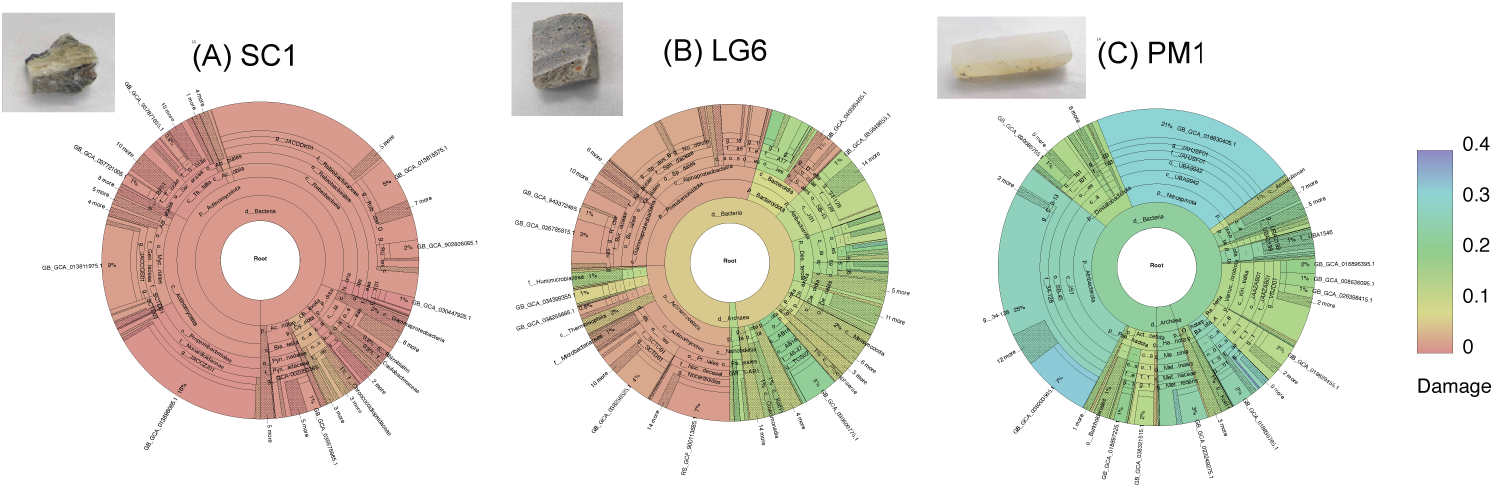
Examples of damage-coloured Krona plots. Plots for all samples are available interactively at https://bdesanctis.github.io/subglacial-precipitates/kronas.html. Each of the 3 circles represents the metagenomic composition of a single subglacial precipitate subsample. Colour is by postmortem damage (5’ C-to-T frequency).

Ordering the most abundant (*>*40,000 total reads) genera across all subglacial precipitate samples by increasing median damage partitions taxa into two distinct sets: ancient, high-damage subglacial taxa, many of which have been found in previous direct subglacial samples, and modern, low-damage surface taxa, almost all of which are found in our Antarctic surface control (Figure 3). We designate taxa to the right of the vertical line in Figure 3 as subglacial. This designation is supported by their consistently high degree of damage, abundance across subglacial precipitate samples, ecotypes, absence from the surface control, presence in previous subglacial records, and in many cases, their complete or near-complete absence from the entire Arctic and Antarctic surface record, including in surface endolithic microbiomes [31, 32, 33]. This separation by damage holds at higher taxonomic orders (Supplementary Figure 3), and with a lower threshold on read count (Supplementary Figures 4-5). Postmortem damage is an effective separator here partly due to the strong taxonomic differences between the ancient subglacial and modern surface microbiomes, and relatively high read counts. This is not always the case - in many systems, the same microbial taxa can persist throughout time, confounding damage signals [34].

**Figure 3:**
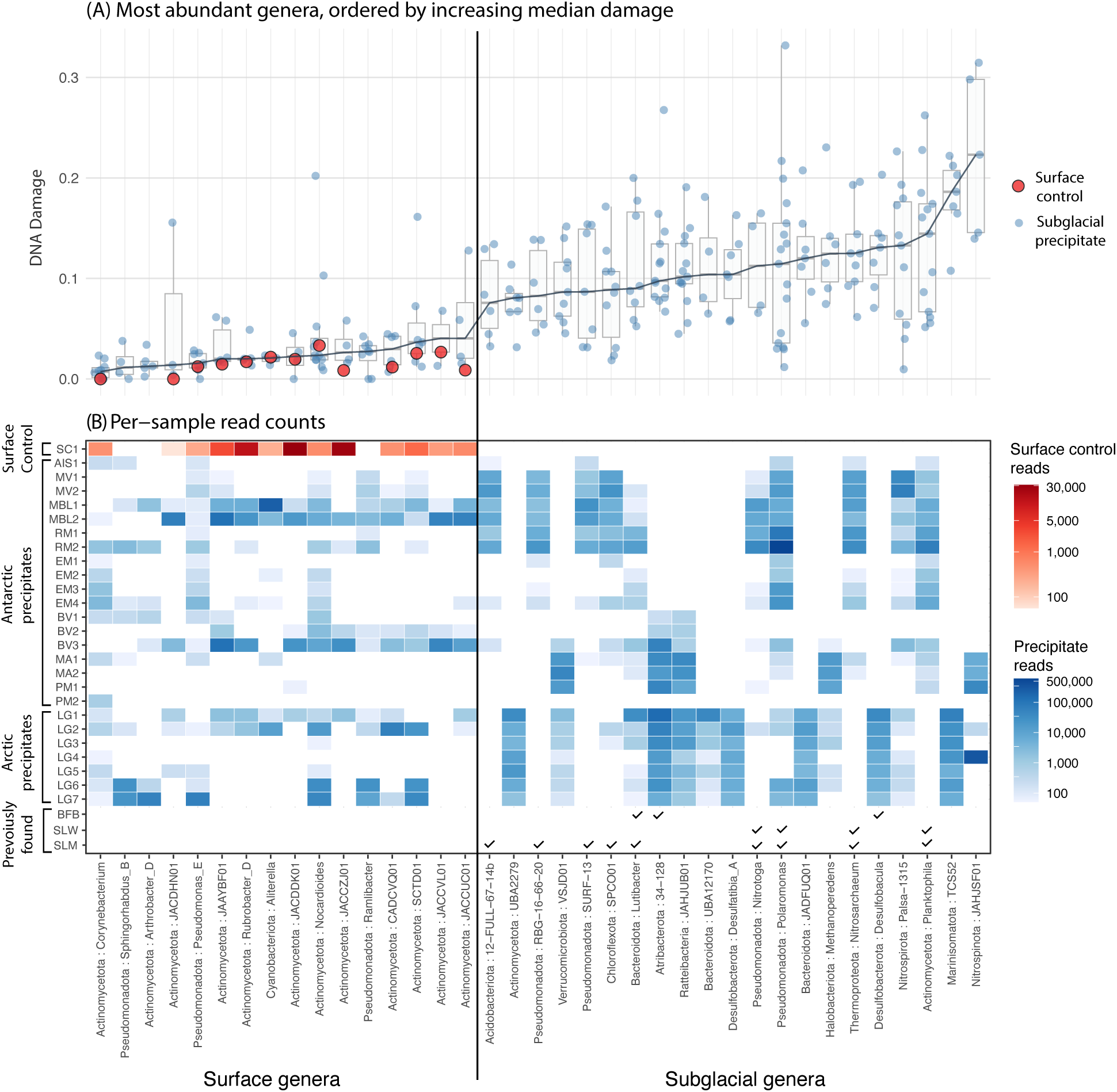
Postmortem damage separates out ancient subglacial taxa from modern surface taxa. (A) All genera with more than 40,000 total reads and present in at least 3 precipitate samples, ordered by increasing median damage (5’ C-to-T) across samples. Precipitate points (blue) are only shown if there were at least 200 reads for the genus in that sample, whereas surface control points (red) are shown if there were at least 50 reads. Genera names are at the bottom, prefixed by their phylum name. (B) Read abundances for the same genera as A, in the same order. Checkmarks are shown if subglacial taxa were previously found via direct sampling in Blood Falls Brine (BFB), Subglacial Lake Whillans (SLW) or Subglacial Lake Mercer (SLM) (see Supplementary Section 5.1).

## 3 Characterizing the subglacial microbiome

Many subglacial taxa we detect with high read counts have previously been reported in modern subglacial environments, link to deep subsurface or extreme cold environments, or have ultra-small genomes or cell sizes. Within several subglacial genera in Figure 3 (*12-FULL-67-14b*, *RBG-16-66-20*, *SURF-13* and *SPCO01*), the top reference, or the reference genome with the most uniquely assigned reads, is a single-cell genome recently isolated from SLM (Subglacial Lake Mercer) [11]. In this SLM study, *SURF-13* was only found in the water column, whereas *SPCO01* was only in sediments, suggesting that our subglacial precipitates are capturing a mixed water and sediment signal. Within Nitrospinota, Ratteibacteria, and Humimicrobiaceae, nearly all reads map uniquely to metagenome-assembled genomes (MAGs) from the Fennoscandian Shield deep groundwater database [35]. The top reference within *Methanoperedens*, a strictly anaerobic methanotrophic archaea, is a MAG assembled from a deep-sea hydrothermal vent [36]. Almost all reads in the subglacial phylum Marinisomatota, a widespread marine heterotroph, are assigned to a single MAG from a cold seep [37]. Within the genus *Lutibacter*, which was previously found in BFB [13], the top reference is a MAG from a hypersaline, anoxic, sub-zero Arctic spring [38]. Most reads within the subglacial genus *UBA12170* only map to a MAG from high Arctic cold saline sulfur spring anaerobic sediment [39]. Anaerolineae is a chemoheterotroph often occurring in marine sediment and also found in SLM [10]. We find two distinct classes of Desulfobacterota, strictly anaerobic sulfate-reducers often associated with deep-sea hydrothermal vents, and previously found in an anoxic, hypersaline Arctic spring [40, 39]. *Nitrotoga*, a cold-tolerant nitrite oxidizer [41] found here, was also detected in high abundance in SLW and SLM [11, 9]. Both *Nitrosarchaeum* and Nanobdellota are in the DPANN superphylum of ultra-small archaea with limited metabolic capabilities, and were previously found in subglacial Antarctica, in SLM/SLW and BFB respectively [42, 11, 9, 13]. Ultrasmall subglacial bacteria identified here include Omnitrophota, Patescibacteria, and *Planktophila*, the latter two of which were previously found in SLW, SLM or BFB [11, 9, 13]. Omnitrophota are chemolithoautotrophs found in anoxic environments such as deep marine sediments or hydrothermal vents [43, 44]. This ultra-small phenotype likely supports efficient or streamlined and potentially interdependent metabolisms in nutrient-limited environments [45].

The subglacial taxa identified here form an abundant and confident subset of the subglacial microbiome, but do not encompass all subglacial microbial diversity, either in these samples or in subglacial waters more broadly. For instance, we can identify further subglacial taxa by lowering our threshold to a minimum of 10,000 total reads (Supplementary Figures 4-5). At this lower threshold, we detect Iainarchaeota, within the archaea superphylum DPANN, for which the top reference is a MAG reconstructed from a deep-sea hydrothermal vent [46]. Similarly we can detect Deferrimicrobia, Ignavibacteria, Terriglobia, CAIJMQ01, JAHJDO01, Aminicenantia, and Desulfitobacteriia (Supplementary Figures 4-5). Some taxa may also inhabit both surface and subglacial environments, potentially leading to highly variable or intermediate damage levels. While this signal is difficult to interpret without prior knowledge, a likely example of this is the Polaromonas genus, which was detected in SLM and SLW [11], is abundant both globally and on glacier surfaces [47], and has highly variable damage values in our samples (Figure 3A).

Although functional analyses and assembly are more difficult to resolve with ancient DNA, we find key functional genes for central metabolic pathways consistent with conditions in directly sampled subglacial environments (Supplementary Section 6-7). For example, genes associated with one-carbon metabolism, including those involved in CO oxidation (cooS/acsA), are present. These genes encode subunits of the carbon monoxide dehydrogenase/acetyl-CoA synthase complex, a core component of the Wood–Ljungdahl pathway of carbon fixation. This pathway is well-suited to subglacial settings because of its low ATP requirements [48] and has been detected in other subglacial environments [49]. We also assembled an early-branching, ancient subglacial Nitrospinota MAG from LG4, which contained genes for near complete denitrification, including the nitrate reduction, nitrite reduction, and nitric oxide reduction steps (Supplementary Section 7). Our data indicate several active biogeochemical processes, including nitrogen and sulfur metabolism and chemosynthetic carbon fixation. Such pathways are essential for subglacial ecosystems, where sunlight is absent and primary production depends on the oxidation of inorganic substrates to fix carbon, generating organic matter for heterotrophic community members [8, 13, 49].

## 4 Subglacial waters partition into two redox states

### 4.1 Partitioning ancient subglacial microbial microbiomes

To compare subglacial microbiomes across precipitate samples, we extracted reads assigned within the most abundant subglacial classes (16 bacterial and 3 archaeal; see Supplementary Figure 3) and performed nonparametric multidimensional scaling on these subglacial taxonomic abundances. This yielded two well-separated clusters (Figure 4A-B). Using OxyMetaG [50], we were able to estimate the relative proportion of aerobic versus anaerobic subglacial bacteria for 17 total samples. Predictions for bacteria in the left cluster samples to range from 58% to 100% aerobic, whereas the right-cluster samples were predicted to contain only anaerobic bacteria (Figure 4C). This implies that oxygen availability is a major driver of subglacial microbiome structure. We therefore refer to the clusters as ‘oxidized’ and ‘reduced’.

**Figure 4:**
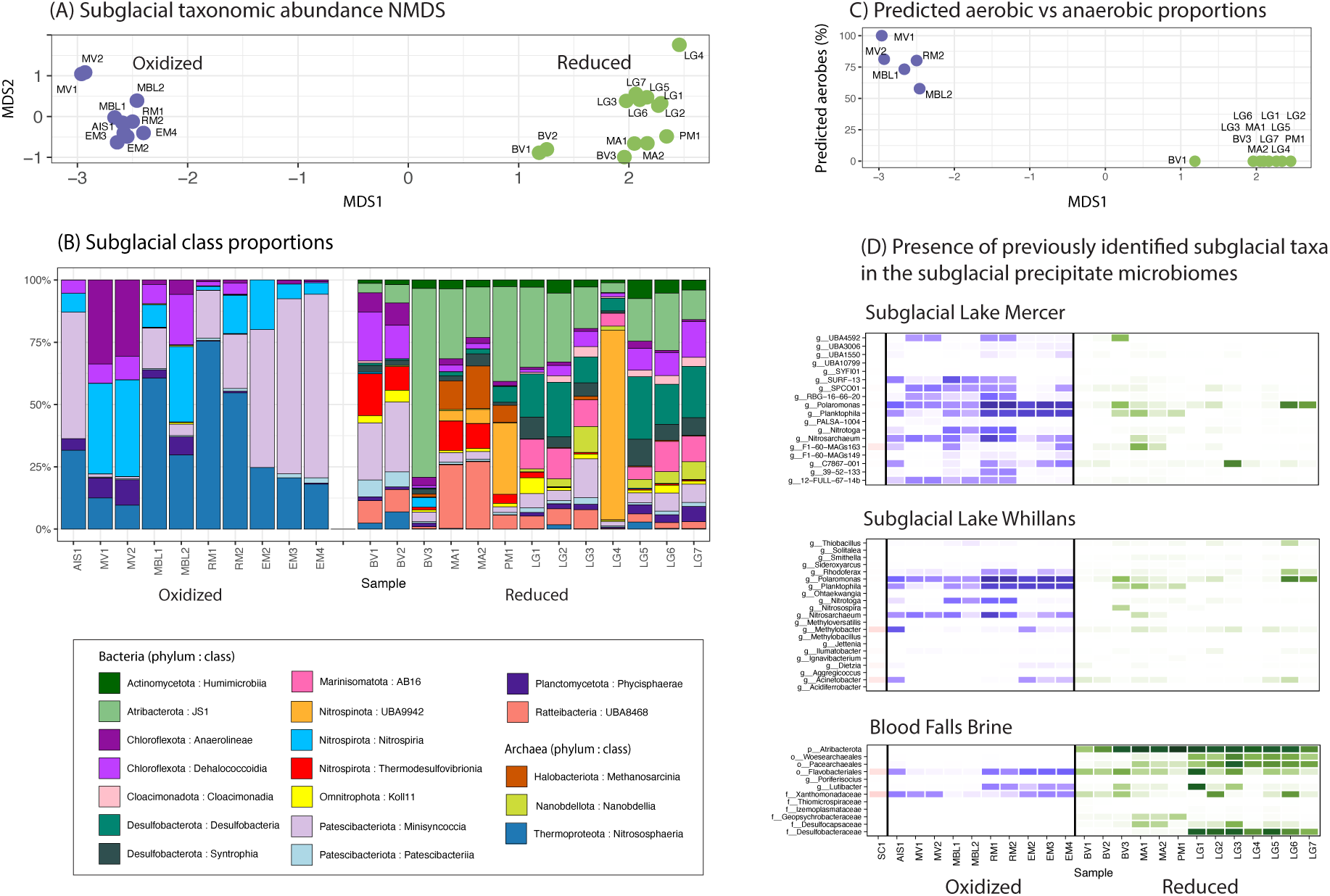
The subglacial microbiome splits into two clusters. (A) An NMDS on the unaggregated read counts of all taxonomic nodes within 19 abundant subglacial classes splits samples into two clear clusters. (B) Proportions of each subglacial class in each sample, grouped by cluster. (C) Predicted proportion of aerobic vs anaerobic bacteria for each sample, using reads which mapped to taxa in the 16 subglacial bacterial classes. (D) Existing Antarctic subglacial samples can be assigned to one of our clusters by assessing the prevalence of their reported top taxa in our samples.

To contextualize the only directly sampled sites in Antarctica, Subglacial Lake Mercer (SLM), Subglacial Lake Whillans (SLW) and Blood Falls Brine (BFB), we assessed the prevalence of the top taxa from these sites in each of our samples, shown in Figure 4D (Supplementary Section 5.1). The direct existing subglacial lake water samples from SLW [9] and SLM [11], the latter of which was saturated with oxygen at the time of sampling, both clearly belong to the oxidized cluster, whereas BFB, which was devoid of measurable oxygen, but suboxic, when sampled, belongs to the reduced cluster [51](Fig. 4E). These cluster memberships, especially BFB, become even clearer upon exclusion of taxa found in our surface control (Figure 4E, SC1), which may be contaminants in the original studies (Supplementary Section 5.1).

The difference between these subglacial clusters is evident in the ancient microbial composition of the precipitates (Figure 4B). Precipitates in the oxidized cluster are rich in *Nitrosarchaeum* (within the Nitrosphaeria class) [11]. *Nitrosarchaeum* has been found in oxic surface layer of marine sediment [42]. The oxidized cluster is abundant in the Nitrospiria class, which was recently found in aerobic groundwater [52]. The reduced cluster is lacking in both Nitrospiria and Nitrosphaeria, but is defined by much more microbial diversity overall. It is notably rich in Atribacterota, particularly class J1 (genus 34 128), which was also found in high abundance in BFB [13]. Atribacterota is a bacterium commonly found in anoxic, methane-rich marine sediments and abundant in deep-sea Antarctic sediment [53]. Within the phylum Nitrospirota, reads map to two abundant classes with contrasting metabolic roles: Nitrospiria, in the oxidized cluster, are aerobic nitrite oxidizers previously found in SLW [9], whereas reads in the reduced cluster map to Thermodesulfovibrionia, obligate anaerobes [54]. Even those classes common to both clusters (e.g. Minisyncoccia, Dehalococcoidia and Anaerolineae) strongly split between clusters at lower taxonomic levels, shown by the per-taxa NMDS loadings (Supplementary Section 4). Note that the Baffin Island subglacial samples (labelled LG) cluster within Antarctic subglacial diversity (Figure 4A), in the reduced cluster, indicating that redox state is a stronger driver of subglacial microbiome composition than geography. We see high consistency in taxonomic composition even down to the genus or species level within each cluster (see Figure 3B and Krona plots), with most of the abundant oxidized or reduced classes in Figure 4B found across all samples in their corresponding cluster.

### 4.2 Iron and manganese concentrations in calcite as indicators of redox state

Oxygen depletion should influence subglacial water geochemistry through the sequential reduction of redox-sensitive elements [55, 26, 56]. For instance, iron and manganese form insoluble ferric and manganic oxides at high redox potentials but become soluble Fe^2+^ and Mn^2+^ under reducing conditions [26]. The introduction of oxygen-rich melt derived from the basal melting of meteoric ice in the ice sheet interior elevates Eh (redox potential) [57], favoring oxidation of Fe^2+^ and Mn^2+^ to Fe^3+^ and Mn^4+^ and precipitation of oxide phases, thereby limiting incorporation into calcite [58, 59]. As microbial respiration consumes oxygen and Eh declines, Fe and Mn are sequentially reduced to their soluble divalent states and become available for lattice substitution into calcite. Because the Mn^4+^ → Mn^2+^ reduction occurs at higher Eh than Fe^3+^ → Fe^2+^, Mn becomes mobilized under mildly reducing conditions, whereas Fe reduction requires more strongly reducing environments. Consequently, calcite Fe and Mn compositions can record a sequence of changes in redox, with low Fe–Mn concentrations reflecting oxic conditions, elevated Mn but low Fe indicating early reduction and enrichment in both Fe and Mn signaling more advanced anoxia [58].

Accordingly, we find that calcite concentrations of Fe and Mn in the subglacial precipitates reproduce the oxidized and reduced clusters (Fig. 5). We measured Fe and Mn concentrations on multiple aliquots from each of the subglacial precipitates analyzed in this study (see Methods for details). The concentrations cluster into two groups: a low-Mn group (∼ 10^1^−10^2^ ppm) with relatively low Fe concentrations (∼ 10^1^−10^3^ ppm), and a high-Mn group (∼ 10^3^-10^4^ ppm) with elevated Fe concentrations (∼ 10^3^-10^4^ ppm) (Figure 5, Supplementary Data 1.3). These clusters exactly match those found using the subglacial microbiomes (Figure 4). The bimodal distribution in iron and manganese concentrations reflects the differing solubility of oxidized Mn^4+^ and Fe^3+^. Calcite with low Mn and Fe concentrations likely formed from relatively oxidized waters, whereas the high-Mn, high-Fe cluster is consistent with precipitation from reduced or anoxic waters. BV1 shows high Mn but low Fe, consistent with an intermediate, partially reduced regime (Figure 5), and is also the most intermediate sample in the reduced cluster as measured by its microbial taxonomic composition (Figure 4A). This association of the microbiomes with calcite Mn and Fe concentrations implies that the oxidized microbiomes occurred in fresh, oxygen-rich meltwater, whereas the reduced microbiomes occurred in low-oxygen waters.

**Figure 5:**
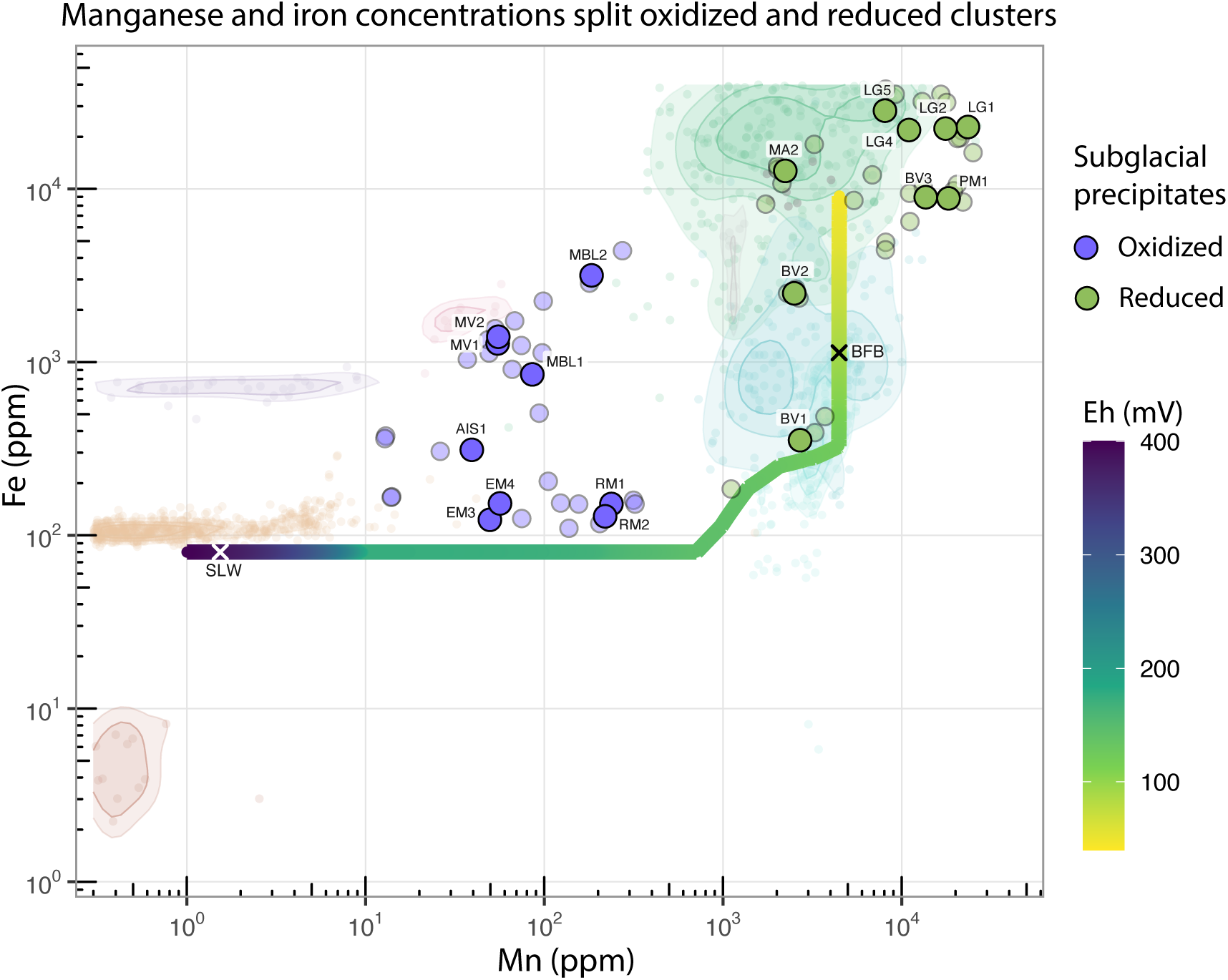
Iron and manganese concentrations in calcite. The Fe and Mn concentration for subglacial precipitates are reported for multiple aliquots (translucent blue and green markers) as well as the mean composition for each sample (opaque, text labeled marker). These are color coded, according to their redox grouping, as defined by taxonomic abundances in Figure 4. A literature compilation of calcite compositions from known redox conditions includes calcite from oxidized settings including speleothems and tufas (warm colored circles/contours) and calcite from reduced settings such as igneous carbonatite and groundwater brine (cool colored circles/contours). Model predicted calcite Mn and Fe concentrations for a range in water Eh modified from ([58]). This permits comparison to modern waters with Eh measurements, including Antarctic subglacial systems with known microbiomes (SLW = Subglacial Lake Whillans; BFB = Blood Falls brine), into the Fe-Mn calcite concentration space.

For comparison, we compiled Fe and Mn concentration data from calcite samples formed under known redox conditions (see Methods and Supplementary Data 1.5-1.10 for data and sources). Speleothems and tufas, which precipitate from oxygenated meteoric waters, are shown as faded background points in warm colors in Figure 5. Like the oxidized subglacial precipitates analyzed here, they contain low concentrations of Mn (*<* 100 ppm) and Fe (*<* 2000 ppm). Calcites formed under reducing conditions, including samples from igneous carbonatites and groundwater brines, are shown as faded background points in cool colors. These show consistently high Mn (*>* 10^3^ ppm) with a wide range of Fe (10^2^–10^5^ ppm), reflecting a redox transition from high Mn, low Fe calcite in partially reduced waters to high Mn, high Fe calcite in anoxic environments. This redox-controlled compositional range matches that observed in the reduced subglacial precipitates.

To refine the relationship between water redox conditions and calcite Fe–Mn concentrations, we refer to the thermodynamic model of Barnaby and Rimstidt [58], which predicts the composition of calcite as a function of water Eh. Here, we adapt their results for batch conditions rather than continuous pore-water evolution (see Methods). The model predicts that high-Eh (oxidizing) waters yield calcite with low Fe and Mn concentrations consistent with oxidized samples in both subglacial precipitates and the literature compilation (Fig 5). As Eh decreases (i.e., with progressive oxygen consumption), Mn and Fe concentrations increase nonlinearly: Mn rises sharply to a maximum (limited by Mn oxide availability) before Fe concentrations begin to increase. This framework enables the mapping of Eh values of natural waters into the Fe-Mn concentration space. In particular, SLW (Eh ≈ 382 mV; [8]) represents oxidized conditions that map onto low-Fe, low-Mn calcite, while BFB (Eh ≈ 90 mV; [60]) corresponds to more reducing or suboxic conditions and calcite enriched in both elements. The agreement between modeled and observed data demonstrates that the model captures the redox control on the iron and manganese composition of calcite, linking calcite geochemistry and direct water oxygen measurements.

## 5 Discussion

Using ancient metagenomics from subglacial precipitates from across Antarctica and the Laurentide ice sheet, we provide a first glimpse of millennia-old subglacial microbiomes. This work expands the spatial and temporal record of the subglacial microbiome to a degree currently unattainable by direct drilling campaigns, and without its associated contamination risks. Our greatly expanded set of subglacial sites includes many taxa and functional genes common in modern subglacial settings. Our samples, along with the 3 cleanly sampled Antarctic sites, partition by oxygen availability into two well-separated clusters, both by microbiome composition and by precipitate geochemistry or redox potential. This work demonstrates the power of ancient metagenomics in reconstructing physically inaccessible environments, beyond its ability to teach us about the past.

Our findings support a framework across poles and ice ages in which subglacial water redox states are held in balance by microbial oxygen consumption and oxygen input from meltwater [1, 3, 4]. Oxygen is introduced to subglacial waters by basal melting of meteoric ice, which releases trapped atmospheric gases [57, 61]. Reduced waters can form when microbial aerobic respiration rates exceed the supply of oxygen from fresh meltwater. Low-oxygen environments will cause microbes to seek out alternate electron acceptors, restructuring the microbiome, and permit the reduction of Mn^4+^ and Fe^3+^ to their soluble Mn^2+^ and Fe^2+^ forms. Thus, reduced precipitates formed in environments with a limited supply of fresh meltwater, whereas oxidized precipitates formed in environments with sufficiently high fresh meltwater input to overcome microbial oxygen respiration rates.

This link between oxygen availability and meltwater flow controls the long-term evolution of subglacial redox conditions and, based on our results, microbial assemblages. Beneath the Antarctic ice sheet, basal meltwater flow is governed by the hydraulic potential gradient, primarily set by ice surface slope [62, 63] and modulated by ice thickness and meltwater input [1]. Southern Hemisphere climate forcing drives changes in Antarctic ice motion and grounding-line position on centennial to millennial timescales [64, 15, 65, 66], and because subglacial hydrology is tightly coupled to ice dynamics [67], this coupling likely regulates basal meltwater flux and oxygen supply. Radioisotopic ages of Antarctic calcite precipitates support this connection: with one exception (MBL1), oxidized precipitates formed during climatically warm intervals defined by Antarctic Dome C *δ*^2^*H* values above −430 ‰[68], whereas reduced samples, with one exception (PM1), date to the last glacial (18 to 55 ka) (Supplementary Fig. 9). Although reduced and oxidized environments can occur in both climate states, as shown by the presence of oxygen-poor brines such as Blood Falls and Lake Vanda [13, 69] under modern warm conditions, the dominant trend in our precipitate data suggests that climate-driven variations in meltwater supply may control oxygen availability, microbial structure, and subglacial water chemistry at sites of precipitate formation.

Reconstructing redox conditions of subglacial waters can inform ocean and climate models by enabling the inclusion of their chemical and biological trajectories. Reduced subglacial environments are more likely to accelerate mineral dissolution and nutrient release from bedrock while promoting the production of climate-reactive gases such as nitrous oxide and methane, which may eventually be discharged to the atmosphere or ocean. The recent emergence of methane seeps in the coastal Antarctic environment of the Ross Sea [70] potentially linked to subglacial groundwater, highlights the significance of these processes. Although we find little to no evidence for microbial methanogenesis (methane production) in our samples, we do detect methanotrophic (methane-consuming) taxa (e.g., Methanoperedens, Methyloceanibacter; see Krona plots) at several locations, an observation consistent with the presence of methanotrophic functional genes and measurable methane-oxidation in modern subglacial lakes [11, 71]. While a potentially vast reservoir of methane may exist beneath the Antarctic ice sheet [3], under favorable redox conditions, the Antarctic subglacial microbiome may function as a methane sink [71]. Understanding microbially driven subglacial redox trends is essential for predicting how these systems shape the solute composition of subglacial groundwater discharge which has potentially far-reaching effects on downstream ecosystem function across space and time.

While direct-access drilling remains essential, the current study could be scaled up substantially using archival material and with a smaller logistical footprint. Individual stratified subglacial precipitates can record cyclical transitions between reduced and oxidized waters that span durations of 50-100 thousand years [15]. Future work might thus aim to characterize the subglacial environment and microbiomes at a single formation site over a 10^3^-10^5^ year time range, providing a closer view into how microbiomes respond to changes in water chemistry and environment. With enough samples, one could build spatiotemporal maps of Antarctic subglacial redox and biogeochemical conditions and hydrological connectivity throughout the Pleistocene, providing constraints for paleomodels of subglacial hydrology and improving predictions of ice sheet behavior and subglacial greenhouse gas emissions under future global warming.

Beyond their role in Earth’s climate and biogeochemical cycles, subglacial environments serve as analogs for some of the most promising targets in the search for extraterrestrial life. Jupiter’s moon Europa, Saturn’s moon Enceladus, and potentially Mars all harbor liquid water beneath or within ice, where life might persist under similar constraints of darkness, isolation, and low temperatures. Identifying environmental factors that fundamentally structure microbial communities in these systems is critical for predicting where habitable niches might exist on icy worlds and what biosignatures might be detectable. Additionally, this work may help select key sampling locales for life-detection missions. For instance, Mars currently hosts two large polar ice caps and numerous glacial landforms, including evidence for extensive deglaciated regions in the southern hemisphere [72]. Radar observations further indicate that the planet experienced periods of widespread glaciation in the past [73, 74, 75], and spectral analyses points to the presence of hydrated silica-bearing deposits [76] that have been interpreted by some as subglacial in origin [77]. Targeting currently and formerly glaciated regions in search of subglacially formed precipitates may be a promising strategy for future Mars missions.

## 6 Methods

### 6.1 Sample collection, dating, and geochemistry

**Subglacial precipitate dating.** The ^234^U-^230^Th age estimates of carbonate formation were obtained at the University of California Santa Cruz (UCSC) Keck Isotope Laboratory from powdered material for all samples following methodology described in [15]. The aDNA studied here is interpreted to be present within subglacial waters and incorporated into the opal or calcite samples during crystallization. The timing of precipitate formation is provided by ^234^U-^230^Th ages, which record the time since the disequilibrium that occurs in surface waters between the radioactive parent ^234^U, which is soluble in waters, and the intermediate daughter ^230^Th, which is insoluble. The calculated dates reflect the ingrowth of ^230^Th back towards secular equilibrium. The analytical methods and isotopic data used to calculate the ages for the samples studied here are reported in references [14, 24, 25, 15, 26].

**Subglacial precipitate geochemistry.** In addition to the timing for formation, the isotopic and chemical compositions of the precipitates can provide insights into the subglacial setting and hydrologic processes, including the chemistry, provenance, redox and salinity of these waters [15, 25]. For example, the oxygen (*δ*^18^O), carbon (*δ*^13^C), and uranium (^234^U/^238^U) compositions can be effective tracers to delineate subglacial waters from other sources, such as surface waters. The oxygen compositions of Antarctic calcite precipitates are some of the most ^18^O-depleted compositions on Earth (−30 to −60 %) [14, 78, 15], matching the compositions of ice found at the interior of the Antarctic ice sheet [79, 80].

The most depleted samples are those samples collected at intra ice moraines (EM, RM, PM, MV), which are up to 15 % more depleted than the local ice, supporting origination from subglacial melting of ice beneath the ice sheet interior [15, 25]. Heavier precipitate samples are those samples found at the ice sheet periphery (BV, MBL) in areas where ice retreat has exposed precipitate samples found on bedrock. The oxygen composition of these samples are still up to 10 % more depleted than local precipitation [79, 80], suggesting a more distal provenance of subglacial waters that migrate through the subglacial hydrologic network to the site of sample precipitation.

The *δ*^13^C of subglacial precipitates exhibit a range of compositions that depends on whether the sample was exhumed at interior moraines (depicted in Figure 1) or formed at the ice sheet periphery and found on bedrock surfaces after Holocene ice retreat. The *δ*^13^C of all moraine samples are highly depleted (−23 to −17 %), suggesting that the dissolved inorganic carbon (DIC) of these subglacial waters was sourced primarily from sedimentary organic matter (−21 to −27 %) and was isolated from the atmosphere. Peripheral samples, are heavier (−8 to −0 %) and when combined with other evidences, suggest carbon sourcing from marine carbonate bedrock (∼0 %).

The [^234^U*/*^238^U] of many subglacial waters, where brackets denote activity ratios, are above secular equilibrium ([^234^U*/*^238^U] = 2-5), an enrichment in [^234^U] that reflects prolonged rock water contact in a subglacial hydrologic system and are matched by measurements of subglacial brines that emit from beneath glaciers at the ice sheet perimeter [81, 82]. In contrast, fresh glacial meltwater exhibits seawater compositions (∼1.14, [83]). In all but one sample site (MBL), subglacial precipitates exhibit elevated [^234^U*/*^238^U] consistent with a subglacial origin, and formation from waters that have experienced prolonged rock water contact.

**Surface control.** We can utilize the oxygen, carbon and uranium isotopic composition from a calcite sample that formed from Antarctic surface melts to illustrate how subglacial waters differ. This sample serves as a surface control for aDNA results. This sample formed a thin 2-3 mm encrustation on the rock outcropping of West Antarctica’s Vivian Nunatak (−77.53°, −143.567°, 21VN01). The carbon *δ*^13^C = −4 to −1 ‰), oxygen *δ*^18^O = −17 to −27 %) and uranium ([^234^U*/*^238^U] = 0.7-1) compositions were measured using the same methods described above. The *δ*^13^C compositions are similar to calcite crusts that form in near equilibrium with the atmosphere [84, 85] implying subaerial formation. The *δ*^18^O are precipitation are heavier than local ice, and correlate *δ*^13^C values, consistent with Arctic carbonates that form by evaporation [84]. The low [^234^U*/*^238^U] do not support long term rock water contact. This calcite, like other polar carbonates [84], is interpreted to form by evaporation of local snow melt.

**Fe-Mn calcite methods.** The major and minor element composition of calcite precipitates were measured using the Thermo iCap 7400 Inductively Coupled Plasma Optical Emission Spectroscopy (ICP-OES) housed in the Plasma Analytical facility at the University of California Santa Cruz. Samples were analyzed for major (Na, K, Mg, Ca), minor (Si, Sr) and trace (P, Fe, Mn) element concentrations. Approximately 10 mg of calcite samples were dissolved in 2% HNO_3_. All samples were spiked internally with yttrium (Y) with concentrations determined using standards at varying concentrations (e.g. concentration calibration curve) formulated by comparing Y-normalized intensities of these standards against the known standard concentrations. Calcite elemental compositions from this effort are reported in the Supplementary Data 1.3 and in Figure 5 where they are compared to a literature compilation of Fe and Mn concentration measurements from calcites that formed from a known redox environment.

The methods used to measure Fe and Mn in speleothems were by LA-ICPMS. The goal of this comparison is to measure the Fe and Mn in the calcite. This method, however, ablates all materials, including any siliciclastics. This biases these measurements, producing high Fe and Mn measurements (when Si is elevated) that follows a steep line in Fe and Mn space, which is the crustal Fe/Mn composition. Thus, we have filtered these data, removing measurements with silicate concentrations above baseline values.

**Modeling calcite Fe and Mn concentrations from water Eh.** To compare measured calcite Fe and Mn concentrations with expected values across a continuum of redox states, we adapted the theoretical thermodynamic framework proposed in [58] for real data. The model describes the continuous evolution of a closed pore-water system in which progressive oxygen consumption drives sequential reduction of Mn(IV) and Fe(III) oxides, resulting in temporally varying aqueous Mn^2+^ and Fe^2+^ activities and corresponding calcite compositions. In the present study, this temporal model was reformulated into a point-Eh equilibrium model that predicts calcite Fe and Mn concentrations for any fixed redox potential (Eh). Each Eh value is treated as a discrete equilibrium state representing a distinct water composition rather than continuous evolution. The resulting relationship defines a non-linear increase in Mn at relatively high Eh, followed by Fe enrichment at lower Eh, reproducing the characteristic Mn-first, Fe-later sequence observed during reduction. We set the model baseline empirically, to the median measured concentrations in a large set of published natural and laboratory calcites with known Fe, Mn and Eh values. This accounts for unavoidable analytical backgrounds, minor lattice substitutions, and geologic variability that produce measurable Fe and Mn even in fully oxidized samples. The model therefore provides a direct mapping between water redox potential and expected calcite Fe–Mn composition, enabling quantitative comparison among subglacial precipitates, literature datasets, and modern subglacial water chemistries of known Eh. This adapted [58] model reproduces the key features observed in both the subglacial and literature datasets. The modeled trend defines a low-Fe, low-Mn field at high Eh (oxidized conditions), a transitional region with elevated Mn and low Fe, and a fully reduced field with enrichment in both elements. The measured calcite compositions from this study plot consistently along this progression. Oxidized subglacial precipitates fall within the model’s high-Eh domain, while samples containing elevated Mn and Fe align with the reduced field. This correspondence is further supported by independent redox measurements from modern Antarctic subglacial environments. The measured Eh of Lake Whillans (≈382 mV; [8]) plots within the oxidized field and coincides with the Fe–Mn compositions of the most oxidized subglacial calcites. In contrast, the Blood Falls subglacial brine (≈90 mV; [60]) occupies the model’s reduced domain and predicts calcite with Fe and Mn concentrations comparable to those of the reduced precipitates analyzed here.

### 6.2 Ancient metagenomic laboratory methods

Sedimentary ancient DNA analyses were carried out on fragments of calcite, opal, or a mix of the two, isolated from individual layers, with total masses ranging from 10^2^ to 10^3^ mg, depending on availability. We performed all extractions in a dedicated ancient DNA clean laboratory at UC Santa Cruz. We wore full PPE and sanitized surfaces with a bleach solution and ethanol before and after working with these samples. Before powdering, we performed a 30 second 0.05% bleach treatment on all samples, followed by three rinses with molecular-grade ultrapure water. In addition, we processed replicates of four samples (MA113, 19ACAG81, Magnis Valley 2, and PRR52588) without bleach-treating to evaluate the effectiveness of bleach treatment on recovery of endogenous DNA (see Supplementary Section 1 and Supplementary Data 1.1 for details). We allowed bleach-treated samples to dry overnight in a dead-air hood in the ancient DNA laboratory. We powdered the dry samples using a dental mallet and bone crusher. One sample (PRR50489 or EM1) was not sufficiently powdered with the bone crusher, so we finished powdering it with a ball mill, which reduced our yield for this sample to 5 mg. We stored all powdered samples in 2 mL screw-cap tubes. Between powdering samples, we cleaned all tools with a bleach solution, rinsed them with ethanol, and irradiated them in a UV crosslinker.

We extracted the powdered samples according to [86] with a 50 mg input for all samples except PRR50489, which only yielded 5 mg after powdering. We extracted in three batches, with one extraction control per batch. We used binding buffer D and high-volume silica spin columns (Roche). We eluted in 50 ul of EBT buffer and quantified the amount of DNA in the extractions using a Qubit with 10 ul sample input. We prepared sequencing libraries using a single-stranded protocol optimized for ancient and degraded DNA [87]. We pooled all libraries equimolarly and sent them to Fulgent Genomics for sequencing to an average depth of 12M reads per sample on a HiSeq (2 x 150bp) instrument (Illumina).

### 6.3 Bioinformatics methods

**Quality control.** We sequenced a total of 37 libraries, including 7 negative controls and four pairs of libraries which were later merged (see Supplementary Methods for details). This yielded a total 1.12 billion raw reads. For each library, we used fastp [88] to merge paired-end reads, remove adapters (using a custom text file containing standard Illumina adapters and residues), remove poly-G and poly-X tails and duplicates, low complexity filter, and discarded reads with low complexity, an average base quality less than 25, or with length less than 30, and retained only merged reads. Duplication rates calculated by fastp were less than 2% in all non-control libraries. We used SGA with a dust threshold of 30, the ropebwt algorithm, and no kmer check to remove low complexity reads [89]. We ran FastQC [90] on each QCed library, and found that base quality declined dramatically after ∼ 150bp into the reads. We therefore cut all filtered merged reads down to 150bp in length.

Using bowtie2 v2.4.4 [91] with default parameters, we first mapped all controls and samples against the human, pig, chicken and cow reference genomes (Accession IDs: GCF 000003025.6, GCF 016699485.2, GCF 002263795, and GCF 000001405.40, [92]), since in our experience these are the most common eukaryotic contaminants in shotgun sequenced ancient environmental DNA. For downstream analysis we removed any reads that mapped to any of these contaminant reference genomes using samtools [93]. The negative controls retained an average of 62% of their reads through this, whereas the surface control and samples retained an average of 94.62% of their reads, with the majority retaining more than 99%.

**Taxonomic profiling.** Next, we downloaded every bacterial and archaeal species representative reference genome in the GTDB database v 226.0 (release226/226.0/genomic files reps/gtdb genomes reps r226.tar.gz, released 2025-04-16 and accessed 2025-06-25), which contains representative microbial genomes and a standardised taxonomy, along with the associated metadata [27]. We concatenated all 143,614 representative species genomes into 15 reference fasta files totalling 455GB uncompressed, and used gtdb to taxdump [94] to generate taxonomy files required for downstream analyses. Both samples and controls were mapped with bowtie2 [91], keeping up to 100 alignments per read per reference file (−k 100), allowing up to 1 mismatch in the seed (−N 1), and default parameters otherwise. Obtaining pairwise alignments is necessary to quantify DNA postmortem damage patterns. Mapping against large reference databases is a common strategy in ancient eukaryotic metagenomic studies and has also been shown to be more sensitive – and often more accurate – than classifier methods [95]. We then used samtools [93] to remove irrelevant reference names in the header in the bam files to which no reads mapped, to merge bams across database pieces, and to query-sort merged bams. Reads were individually assigned to taxonomic nodes using the lowest-common-ancestor approach ngsLCA [95] using a minimum similarity of 95% for all samples and controls. This mapping and lowestcommon-ancestor approach is a common strategy in ancient eukaryotic metagenomics that has successfully been extended to ancient microbial metagenomics [96, 97].

In order to stringently decontaminate our samples, we first assessed the taxonomic composition of 7 negative controls (extraction and library blanks). To do this, we ran bamdam [98] shrink and compute on the negative controls in single-stranded mode with a minimum read count of 10, up to a taxonomic level of “phylum”, a minimum similarity of 95%, and default parameters otherwise. Across the negative controls this yielded 930 unique taxonomic IDs with more than 50 reads in any control. Genera with more than 1000 assigned reads in at least one negative control sample were the human or skin associated bacteria *Cutibacterium*, *Staphylococcus*, and *Corynebacterium*, lab contaminants *Bradyrhizobium*, *Sphingomonas*, *Acinetobacter*, *Leuconostoc*, *Xanthomonas*, and *Pseudomonas*, and lastly *Calothrix*, a freshwater cyanobacteria. For the most part these are well-known contaminants [99]. All negative controls showed fairly similar taxonomic profiles, although only negative control library was abundant in *Calothrix*.

We next ran bamdam [98] on the bam files and ngsLCA output from the 30 samples, including the surface control, in single-stranded mode, a taxonomic level of “phylum” or below, a minimum similarity of 95%, and excluded all reads which were assigned to any of the 930 taxonomic nodes identified with more than 50 reads in any of the controls, and default parameters otherwise. Excluding reads assigned to all taxonomic nodes identified with more than 50 reads in the negative controls is a stringent filter; the most abundant negative control contaminants identified had tens of thousands of reads assigned in the controls, as low as the subspecies level. Exclusion in bamdam is performed on a per-node basis: we did not necessarily exclude all taxonomic nodes *underneath* those identified in the controls; for example, if a phylum appeared in the controls, we did not remove all of the species underneath that phylum (unless we also saw those species in the controls). Next, we examined the bamdam output and found that the taxonomic composition of the 4 samples which were experimental replicates (in two cases, each with one library with bleach treatment and one without) was very similar between replicates. As such, we merged the bam and lca files for each pair of replicates, and re-ran bamdam [98] (with the same parameters) on the merged files, leaving us with 26 total unique samples (excluding controls). We kept separate samples which were from different sections of the same subglacial precipitate, such as MA1 and MA2, which were from different parts of precipitate MA113 (see Supplementary Methods). We also ran bamdam on all 26 samples separately without removing the control taxa to determine the effect of excluding taxa in this way. As can be seen in Supplementary Figure 1, this decontamination removes a minority of taxa and reads for most samples. There were a total of 9.53 million reads assigned to phylum level or below (see Krona plots, Supplementary Section 2 and Supplementary Figure 2), across both the negative controls and the decontaminated samples and surface control used going forward, more than enough for accurate reference-based taxonomic analysis [100]. Damage-coloured krona plots were created with bamdam krona [98] and KronaTools ktImportXML [30].

At this point we also ran the metagenomics profiler sylph v0.6.1 [101] on all samples and controls (after removing reads mapped to human, chicken pig and cow as above) with default options and the GTDB database to compare to our alignment-based results. Results from sylph were a strict subset of alignment-based results, yielding only the microbial taxa already seen in the mapping results with the most reads in each sample, averaging only 8.7 taxa per sample with a maximum of 35.

We transformed the bamdam output files into abundance and damage matrices using bamdam combine, with one row per taxonomic node. After analyzing taxonomic results, read counts and damage values (e.g. Figure 3, Supplementary Figures 3-5), we decided to use class as a taxonomic cutoff for subsequent analyses. 19 subglacial classes (16 bacteria, 3 archaea) were designated according to Supplementary Figure 3 (Figure 4B). We extracted only these taxonomic nodes within these subglacial classes from the bamdam tsv files, and ran an NMDS on the unaggregated read counts for the 4390 subglacial taxonomic nodes using metaMDS from the R vegan package [102] (Figure 4A), on only those samples with more than 200 reads (discarding EM1, PM1, SC1, and all negative controls). Top taxa were also pulled from publications regarding the three Antarctic subglacial sites SLM, SLW and BFB, as detailed in Supplementary Section 5.1.

**Oxygen metabolism**. We used OxyMetaG [50] to predict the relative abundance of aerobic and anaerobic bacteria. OxyMetaG is a ratio-based, gene-based method that was built using phenotypic data from BacDive [103], trained on simulated metagenomes with known proportions of anaerobes and aerobes, and validated on a suite of modern metagenomes with known oxygen levels. We used our read-mapping based pipeline to extract just subglacial bacterial reads for each sample (reads assigned to taxonomic nodes within the 19 high-abundance subglacial classes), which were used as inputs to OxyMetaG. We used ‘oxymetag profile’ to calculate the abundances of 20 marker genes of bacterial oxygen tolerance (13 aerobic markers, 7 anaerobic markers). We then used ‘oxymetag predict’ with ‘-m custom’ to test several hit cutoffs and calculate the ratio of aerobic marker genes to anaerobic marker genes and predict the percent aerobes using the generalized additive model established with the simulated metagenomes. We present results using relaxed cutoffs of percent identity ≥ 45%, e-value *<* 0.1, and bitscore ≥ 25, but we note that results were similar when using more stringent cutoffs of percent identity ≥ 60% and e-value *<* 0.001. We retained predictions for samples with at least 3 of the marker genes detected. This yielded predictions for 5 oxidized-cluster samples and 12 reduced-cluster samples. The number of subglacial bacterial reads in those samples ranged from 6,127 to 489,925. If both anaerobic and aerobic genes were detected, the abundance ratio was used for prediction. There were 6 instances where ≥ 3 anaerobic genes were detected and 0 aerobic genes were detected, and in this case the predicted percent aerobes was set to 0. There were no instances of ≥ 3 aerobic genes detected and 0 anaerobic genes detected.

R packages used for analysis and plotting include tidyverse [104], rnaturalearth [105], sf [106], and vegan [102]. Computations were done primarily on the UCSC Genomics Institute computing server.

Additional information can be found in the Supplement. Code is available at https://github.com/bdesanctis/subglacial-precipitates/.

## Acknowledgements

This paper is dedicated to the memory of Dr. Sarah Crump, who led the genomic data generation in this study. She was a leader in the field of sedimentary ancient DNA, and a mentor, friend, and inspiration to many.

We thank Anne Nakamoto, Jen Quick-Cleveland, Ruairidh Macleod, Yucheng Wang, and Michael Hoffert for helpful comments and discussions. We are grateful to the UCSC genomics institute computing infrastructure team.

BDS would like to acknowledge support from the President’s Postdoctoral Fellowship Program (PPFP). NBD was supported by NSF 21-567 from the US National Science Foundation Office of Polar Programs and by the Center for Microbial Exploration, University of Colorado Boulder. CW recognizes support from the National Institute of Health under Fellowship T32HG012344. CPB was funded by the Novo Nordisk Foundation. AR was supported by Carl Trygger Stiftelse. This research was funded by NSF 2042495 to T.B., and NSF 2045611 to E.T.R. This research is based on services provided by the Polar Rock Repository with support from the National Science Foundation, under Award OPP-2436582.

## Author Contributions

BDS led the bioinformatics analyses, with help from NBD, CPB, RCD, and JM. NBD performed the functional annotation and MAG assembly. CPB conducted the oxygen metabolism analysis and MAG annotation. JJW conducted the oxygen tolerance analysis on the previously sampled sites. SC led the ancient DNA wet lab work and generated the sequencing data, with help from CW and HB. TB supplied the subglacial precipitate samples and led the dating and geochemistry analyses, with assistance from AH, TR, BN, GE, and GP. BDS, JW, TB, and AR wrote the paper, with input from all authors.

## Data availability

Raw sequencing data will be deposited in the European Nucleotide Archive (ENA) and made publicly available upon publication. All other data generated in this study are available in Supplementary Data 1.1-1.10.

## Notes

### Competing Interest Statement

The authors have declared no competing interest.

## References

[1] Stephen J. Livingstone, Yan Li, Anja Rutishauser, Rebecca J. Sanderson, Kate Winter, Jill A. Mikucki, Helgi Bjornsson, Jade S. Bowling, Winnie Chu, Christine F. Dow, Helen A. Fricker, Malcolm McMillan, Felix S. L. Ng, Neil Ross, Martin J. Siegert, Matthew Siegfried, and Andrew J. Sole. Subglacial lakes and their changing role in a warming climate. Nature Reviews Earth & Environment, 3(2):106–124, January 2022.

[2] Mark J. Hopwood, Dustin Carroll, Thorben Dunse, Andy Hodson, Johnna M. Holding, Jose L. Iriarte, Sofia Ribeiro, Eric P. Achterberg, Carolina Cantoni, Daniel F. Carlson, Melissa Chierici, Jennifer S. Clarke, Stefano Cozzi, Agneta Fransson, Thomas Juul-Pedersen, Mie H. S. Winding, and Lorenz Meire. Review article: How does glacier discharge affect marine biogeochemistry and primary production in the Arctic? The Cryosphere, 14(4):1347–1383, April 2020.

[3] J. L. Wadham, S. Arndt, S. Tulaczyk, M. Stibal, M. Tranter, J. Telling, G. P. Lis, E. Lawson, A. Ridgwell, A. Dubnick, M. J. Sharp, A. M. Anesio, and C. E. H. Butler. Potential methane reservoirs beneath Antarctica. Nature, 488(7413):633–637, August 2012.

[4] Martyn Tranter, Mark Skidmore, and Jemma Wadham. Hydrological controls on microbial communities in subglacial environments. Hydrological Processes, 19(4):995–998, 2005.

[5] Bernd Dachwald, Stephan Ulamec, Frank Postberg, Frank Sohl, Jean-Pierre de Vera, Christoph Waldmann, Ralph D. Lorenz, Kris A. Zacny, Hugo Hellard, Jens Biele, and Petra Rettberg. Key technologies and instrumentation for subsurface exploration of ocean worlds. Space Science Reviews, 216(5), June 2020.

[6] John C. Priscu, Amanda M. Achberger, Joel E. Cahoon, Brent C. Christner, Robert L. Edwards, Warren L. Jones, Alexander B. Michaud, Matthew R. Siegfried, Mark L. Skidmore, Robert H. Spigel, Gregg W. Switzer, Slawek Tulaczyk, and Trista J. Vick-Majors. A microbiologically clean strategy for access to the Whillans Ice Stream subglacial environment. Antarctic Science, 25(5):637–647, March 2013.

[7] J.A. Mikucki, C.G. Schuler, I. Digel, J. Kowalski, M.J. Tuttle, M. Chua, R. Davis, A.M. Purcell, D. Ghosh, G. Francke, M. Feldmann, C. Espe, D. Heinen, B. Dachwald, J. Clemens, W.B. Lyons, and S. Tulaczyk. Field-based planetary protection operations for melt probes: Validation of clean access into the Blood Falls, Antarctica, englacial ecosystem. Astrobiology, 23(11):1165–1178, November 2023.

[8] Brent C. Christner, John C. Priscu, Amanda M. Achberger, Carlo Barbante, Sasha P. Carter, Knut Christianson, Alexander B. Michaud, Jill A. Mikucki, Andrew C. Mitchell, Mark L. Skidmore, Trista J. Vick-Majors, W. P. Adkins, S. Anandakrishnan, G. Barcheck, L. Beem, A. Behar, M. Beitch, R. Bolsey, C. Branecky, R. Edwards, A. Fisher, H. A. Fricker, N. Foley, B. Guthrie, T. Hodson, H. Horgan, R. Jacobel, S. Kelley, K. D. Mankoff, E. McBryan, R. Powell, A. Purcell, D. Sampson, R. Scherer, J. Sherve, M. Siegfried, and S. Tulaczyk. A microbial ecosystem beneath the West Antarctic ice sheet. Nature, 512(7514):310–313, August 2014.

[9] Amanda M. Achberger, Brent C. Christner, Alexander B. Michaud, John C. Priscu, Mark L. Skidmore, and Trista J. Vick-Majors. Microbial community structure of Subglacial Lake Whillans, West Antarctica. Frontiers in Microbiology, 7, September 2016.

[10] Christina L Davis, Ryan A Venturelli, Alexander B Michaud, Jon R Hawkings, Amanda M Achberger, Trista J Vick-Majors, Brad E Rosenheim, John E Dore, August Steigmeyer, Mark L Skidmore, Joel D Barker, Liane G Benning, Matthew R Siegfried, John C Priscu, Brent C Christner, Carlo Barbante, Mark Bowling, Justin Burnett, Timothy Campbell, Billy Collins, Cindy Dean, Dennis Duling, Helen A Fricker, Alan Gagnon, Christopher Gardner, Dar Gibson, Chloe Gustafson, David Harwood, Jonas Kalin, Kathy Kasic, Ok-Sun Kim, Edwin Krula, Amy Leventer, Wei Li, W Berry Lyons, Patrick McGill, James McManis, David McPike, Anatoly Mironov, Molly Patterson, Graham Roberts, James Rot, Cathy Trainor, Martyn Tranter, John Winans, and Bob Zook. Biogeochemical and historical drivers of microbial community composition and structure in sediments from Mercer Subglacial Lake, West Antarctica. ISME Communications, 3(1), January 2023.

[11] Kyung Mo Kim, Kyuin Hwang, Hanbyul Lee, Ahnna Cho, Christina L. Davis, Brent C. Christner, John C. Priscu, and Ok-Sun Kim. Genetic isolation and metabolic complexity of an Antarctic subglacial microbiome. Nature Communications, 16(1), August 2025.

[12] Jill A. Mikucki and John C. Priscu. Bacterial diversity associated with Blood Falls, a subglacial outflow from the Taylor Glacier, Antarctica. Applied and Environmental Microbiology, 73(12):4029–4039, June 2007.

[13] Richard Campen, Julia Kowalski, W. Berry Lyons, Slawek Tulaczyk, Bernd Dachwald, Erin Pettit, Kathleen A. Welch, and Jill A. Mikucki. Microbial diversity of an Antarctic subglacial community and high-resolution replicate sampling inform hydrological connectivity in a polar desert. Environmental Microbiology, 21(7):2290–2306, April 2019.

[14] T. Blackburn, G. H. Edwards, S. Tulaczyk, M. Scudder, G. Piccione, B. Hallet, N. McLean, J. C. Zachos, B. Cheney, and J. T. Babbe. Ice retreat in Wilkes Basin of East Antarctica during a warm interglacial. Nature, 583(7817):554–559, July 2020.

[15] Gavin Piccione, Terrence Blackburn, Slawek Tulaczyk, E. Troy Rasbury, Mathis P. Hain, Daniel E. Ibarra, Katharina Methner, Chloe Tinglof, Brandon Cheney, Paul Northrup, and Kathy Licht. Subglacial precipitates record Antarctic ice sheet response to late Pleistocene millennial climate cycles. Nature Communications, 13(1), September 2022.

[16] Richard Bintanja. On the glaciological, meteorological, and climatological significance of antarctic blue ice areas. Reviews of Geophysics, 37(3):337–359, August 1999.

[17] Christine M. Kassab, Kathy J. Licht, Rickard Petersson, Katrin Lindback, Joseph A. Graly, and Michael R. Kaplan. Formation and evolution of an extensive blue ice moraine in central transantarctic mountains, antarctica. Journal of Glaciology, 66(255):49–60, November 2019.

[18] C. L. Freeman, L. Dieudonne, O. B. A. Agbaje, M. Zure, J. Q. Sanz, M. Collins, and K. K. Sand.̌ Survival of environmental DNA in sediments: Mineralogic control on DNA taphonomy. Environmental DNA, 5(6):1691–1705, November 2023.

[19] K. K. Sand, S. Jelavic, K. H. Kjær, and A. Prohaska. Importance of eDNA taphonomy and sediment provenance for robust ecological inference: Insights from interfacial geochemistry. Environmental DNA, 6(2), March 2024.

[20] Tomas Lindahl. Instability and decay of the primary structure of DNA. Nature, 362(6422):709–715, April 1993.

[21] Elise D. Snyder, Jennifer L. Tank, Abagael N. Pruitt, Brett Peters, Pedro F. P. Brandao-Dias, E. M. Curtis, Kyle Bibby, Arial J. Shogren, Diogo Bolster, Scott P. Egan, and Gary A. Lamberti. Warming increases environmental DNA removal rates in flowing waters. Environmental DNA, 7(3), May 2025.

[22] Anne Marie Høier Eriksen, Juan Antonio Rodrıguez, Frederik Seersholm, Hege Ingjerd Hollund, Anne Birgitte Gotfredsen, Matthew James Collins, Bjarne Grønnow, Mikkel Winther Pedersen, M. Thomas P. Gilbert, and Henning Matthiesen. Exploring DNA degradation in situ and in museum storage through genomics and metagenomics. Communications Biology, 8(1), February 2025.

[23] Mateusz Kciuk, Beata Marciniak, Mariusz Mojzych, and Renata Kontek. Focus on UV-induced DNA damage and repair—disease relevance and protective strategies. International Journal of Molecular Sciences, 21(19):7264, October 2020.

[24] Graham H. Edwards, Terrence Blackburn, Gavin Piccione, Slawek Tulaczyk, Gifford H. Miller, and Cosmo Sikes. Terrestrial evidence for ocean forcing of Heinrich events and subglacial hydrologic connectivity of the Laurentide Ice Sheet. Science Advances, 8(42), October 2022.

[25] Jessica Gagliardi, Terrence Blackburn, Gavin Piccione, Slawek Tulaczyk, and C. Brenhin Keller. Subglacial precipitates record Antarctic Ice Sheet response to Southern Ocean warming. Geophysical Research Letters, 52(8), April 2025.

[26] Gavin Piccione, Terrence Blackburn, Paul Northrup, Slawek Tulaczyk, and Troy Rasbury. Antarctic subglacial trace metal mobility linked to climate change across termination III. The Cryosphere, 19(6):2247–2261, June 2025.

[27] Donovan H Parks, Pierre-Alain Chaumeil, Aaron J Mussig, Christian Rinke, Maria Chuvochina, and Philip Hugenholtz. GTDB release 10: a complete and systematic taxonomy for 715230 bacterial and 17245 archaeal genomes. Nucleic Acids Research, October 2025.

[28] Silvia Frisia, Laura S. Weyrich, John Hellstrom, Andrea Borsato, Nicholas R. Golledge, Alexandre M. Anesio, Petra Bajo, Russell N. Drysdale, Paul C. Augustinus, Camille Rivard, and Alan Cooper. The influence of Antarctic subglacial volcanism on the global iron cycle during the Last Glacial Maximum. Nature Communications, 8(1), June 2017.

[29] Adrian W. Briggs, Udo Stenzel, Philip L. F. Johnson, Richard E. Green, Janet Kelso, Kay Prufer, Matthias Meyer, Johannes Krause, Michael T. Ronan, Michael Lachmann, and Svante Paabo. Patterns of damage in genomic DNA sequences from a Neandertal. Proceedings of the National Academy of Sciences, 104(37):14616–14621, September 2007.

[30] Brian D Ondov, Nicholas H Bergman, and Adam M Phillippy. Interactive metagenomic visualization in a web browser. BMC Bioinformatics, 12(1), September 2011.

[31] Vesselin V. Doytchinov and Svetoslav G. Dimov. Microbial community composition of the Antarctic ecosystems: Review of the bacteria, fungi, and archaea identified through an NGS-based metagenomics approach. Life, 12(6):916, June 2022.

[32] Nicholas B. Dragone, Mary K. Childress, Caihong Vanderburgh, Rachel Willmore, Ian D. Hogg, Leopoldo G. Sancho, Charles K. Lee, John E. Barrett, C. Alisha Quandt, Joshua J. LeMonte, Byron J. Adams, and Noah Fierer. A comprehensive survey of soil microbial diversity across the Antarctic continent. Polar Biology, 48(2), February 2025.

[33] Claudia Coleine, Davide Albanese, Angelique E. Ray, Manuel Delgado-Baquerizo, Jason E. Stajich, Timothy J. Williams, Stefano Larsen, Susannah Tringe, Christa Pennacchio, Belinda C. Ferrari, Claudio Donati, and Laura Selbmann. Metagenomics untangles potential adaptations of Antarctic endolithic bacteria at the fringe of habitability. Science of The Total Environment, 917:170290, March 2024.

[34] Carl-Eric Wegner, Raphaela Stahl, Irina Velsko, Alex Hubner, Zandra Fagernas, Christina Warinner, Robert Lehmann, Thomas Ritschel, Kai U. Totsche, and Kirsten Kusel. A glimpse of the paleome in endolithic microbial communities. Microbiome, 11(1), September 2023.

[35] Maliheh Mehrshad, Margarita Lopez-Fernandez, John Sundh, Emma Bell, Domenico Simone, Moritz Buck, Rizlan Bernier-Latmani, Stefan Bertilsson, and Mark Dopson. Energy efficiency and biological interactions define the core microbiome of deep oligotrophic groundwater. Nature Communications, 12(1), July 2021.

[36] Gregory J. Dick. The microbiomes of deep-sea hydrothermal vents: distributed globally, shaped locally. Nature Reviews Microbiology, 17(5):271–283, March 2019.

[37] Yingchun Han, Chuwen Zhang, Zhuoming Zhao, Yongyi Peng, Jing Liao, Qiuyun Jiang, Qing Liu, Zongze Shao, and Xiyang Dong. A comprehensive genomic catalog from global cold seeps. Scientific Data, 10(1), September 2023.

[38] Elisse Magnuson, Ianina Altshuler, Miguel A Fernandez-Martınez, Ya-Jou Chen, Catherine Maggiori, Jacqueline Goordial, and Lyle G Whyte. Active lithoautotrophic and methane-oxidizing microbial community in an anoxic, sub-zero, and hypersaline high Arctic spring. The ISME Journal, 16(7):1798–1808, April 2022.

[39] Elisse Magnuson, Ianina Altshuler, Nastasia J. Freyria, Richard J. Leveille, and Lyle G. Whyte. Sulfur-cycling chemolithoautotrophic microbial community dominates a cold, anoxic, hypersaline Arctic spring. Microbiome, 11(1), September 2023.

[40] Rashmi Saini, Rupam Kapoor, Rita Kumar, T.O. Siddiqi, and Anil Kumar. Co2 utilizing microbes — a comprehensive review. Biotechnology Advances, 29(6):949–960, November 2011.

[41] Sabine Keuter, Hanna Koch, Katharina Sass, Simone Wegen, Natuschka Lee, Sebastian Lucker, and Eva Spieck. Some like it cold: the cellular organization and physiological limits of cold-tolerant nitriteoxidizing Nitrotoga. Environmental Microbiology, 24(4):2059–2077, March 2022.

[42] Dayu Zou, Yanling Qi, Jinjie Zhou, Yang Liu, and Meng Li. Unveiling the life of archaea in sediments: Diversity, metabolic potentials, and ecological roles. iMetaOmics, 2(1), January 2025.

[43] Jialin Hou, Stefan M. Sievert, Yinzhao Wang, Jeffrey S. Seewald, Vengadesh Perumal Natarajan, Fengping Wang, and Xiang Xiao. Microbial succession during the transition from active to inactive stages of deep-sea hydrothermal vent sulfide chimneys. Microbiome, 8(1), June 2020.

[44] Jana Glockner, Michael Kube, Pravin Malla Shrestha, Marc Weber, Frank Oliver Glockner, Richard Reinhardt, and Werner Liesack. Phylogenetic diversity and metagenomics of candidate division OP3. Environmental Microbiology, 12(5):1218–1229, April 2010.

[45] Rose S. Kantor, Kelly C. Wrighton, Kim M. Handley, Itai Sharon, Laura A. Hug, Cindy J. Castelle, Brian C. Thomas, and Jillian F. Banfield. Small genomes and sparse metabolisms of sediment-associated bacteria from four candidate phyla. mBio, 4(5), November 2013.

[46] John D Carlton, Marguerite V Langwig, Xianzhe Gong, Emily J Aguilar-Pine, Mirna Vazquez-RosasLanda, Kiley W Seitz, Brett J Baker, and Valerie De Anda. Expansion of Armatimonadota through marine sediment sequencing describes two classes with unique ecological roles. ISME Communications, 3(1), June 2023.

[47] Jan Gawor, Jakub Grzesiak, Joanna Sasin-Kurowska, Piotr Borsuk, Robert Gromadka, Dorota Gorniak, Aleksander Swiatecki, Tamara Aleksandrzak-Piekarczyk, and Marek K. Zdanowski. Evidence of adaptation, niche separation and microevolution within the genus Polaromonas on Arctic and Antarctic glacial surfaces. Extremophiles, 20(4):403–413, April 2016.

[48] Arren Bar-Even, Elad Noor, and Ron Milo. A survey of carbon fixation pathways through a quantitative lens. Journal of Experimental Botany, 63(6):2325–2342, December 2011.

[49] J. A. Mikucki, P. A. Lee, D. Ghosh, A. M. Purcell, A. C. Mitchell, K. D. Mankoff, A. T. Fisher, S. Tulaczyk, S. Carter, M. R. Siegfried, H. A. Fricker, T. Hodson, J. Coenen, R. Powell, R. Scherer, T. Vick-Majors, A. A. Achberger, B. C. Christner, and M. Tranter. Subglacial Lake Whillans microbial biogeochemistry: a synthesis of current knowledge. Philosophical Transactions of the Royal Society A: Mathematical, Physical and Engineering Sciences, 374(2059):20140290, January 2016.

[50] C.P. Bueno de Mesquita, E. Stallard-Olivera, and N. Fierer. Predicting the proportion of aerobic and anaerobic bacteria from metagenomic reads with OxyMetaG. GitHub, 2025. GitHub: https://github.com/cliffbueno/oxymetag.

[51] J. A. Mikucki, E. Auken, S. Tulaczyk, R. A. Virginia, C. Schamper, K. I. Sørensen, P. T. Doran, H. Dugan, and N. Foley. Deep groundwater and potential subsurface habitats beneath an Antarctic dry valley. Nature Communications, 6(1), April 2015.

[52] Asher E. Keithley, Hodon Ryu, Vicente Gomez-Alvarez, Stephen Harmon, Christina Bennett-Stamper, Daniel Williams, and Darren A. Lytle. Comprehensive characterization of aerobic groundwater biotreatment media. Water Research, 230:119587, February 2023.

[53] Stephanie A. Carr, Beth N. Orcutt, Kevin W. Mandernack, and John R. Spear. Abundant Atribacteria in deep marine sediment from the Adelie Basin, Antarctica. Frontiers in Microbiology, 6, August 2015.

[54] Holger Daims and Michael Wagner. Nitrospira. Trends in Microbiology, 26(5):462–463, May 2018.

[55] Jon R. Hawkings, Mark L. Skidmore, Jemma L. Wadham, John C. Priscu, Peter L. Morton, Jade E. Hatton, Christopher B. Gardner, Tyler J. Kohler, Marek Stibal, Elizabeth A. Bagshaw, August Steigmeyer, Joel Barker, John E. Dore, W. Berry Lyons, Martyn Tranter, and Robert G. M. Spencer. Enhanced trace element mobilization by earth’s ice sheets. Proceedings of the National Academy of Sciences, 117(50):31648–31659, November 2020.

[56] Stephan Krisch, Mark James Hopwood, Janin Schaffer, Ali Al-Hashem, Juan Hofer, Michiel M. Rutgers van der Loeff, Tim M. Conway, Brent A. Summers, Pablo Lodeiro, Indah Ardiningsih, Tim Steffens, and Eric Pieter Achterberg. The 79°N glacier cavity modulates subglacial iron export to the NE Greenland Shelf. Nature Communications, 12(1), May 2021.

[57] Joseph A. Graly, James I. Drever, and Neil F. Humphrey. Calculating the balance between atmospheric CO2 drawdown and organic carbon oxidation in subglacial hydrochemical systems. Global Biogeochemical Cycles, 31(4):709–727, April 2017.

[58] Roger J. Barnaby and J. Donald Rimstidt. Redox conditions of calcite cementation interpreted from Mn and Fe contents of authigenic calcites. Geological Society of America Bulletin, 101(6):795–804, June 1989.

[59] Edward L. Dromgoole and Lynn M. Walter. Iron and manganese incorporation into calcite: Effects of growth kinetics, temperature and solution chemistry. Chemical Geology, 81(4):311–336, February 1990.

[60] Jill A. Mikucki, Ann Pearson, David T. Johnston, Alexandra V. Turchyn, James Farquhar, Daniel P. Schrag, Ariel D. Anbar, John C. Priscu, and Peter A. Lee. A contemporary microbially maintained subglacial ferrous “ocean”. Science, 324(5925):397–400, April 2009.

[61] Alexander B. Michaud and John C. Priscu. Sediment oxygen consumption in antarctic subglacial environments. Limnology and Oceanography, 68(7):1557–1566, May 2023.

[62] R. L. Shreve. Movement of water in glaciers. Journal of Glaciology, 11(62):205–214, 1972.

[63] Frank Pattyn. Investigating the stability of subglacial lakes with a full stokes ice-sheet model. Journal of Glaciology, 54(185):353–361, 2008.

[64] Daniel P. Lowry, Holly K. Han, Nicholas R. Golledge, Natalya Gomez, Katelyn M. Johnson, and Robert M. McKay. Ocean cavity regime shift reversed west antarctic grounding line retreat in the late holocene. Nature Communications, 15(1), April 2024.

[65] Sarah U. Neuhaus, Slawek M. Tulaczyk, Nathan D. Stansell, Jason J. Coenen, Reed P. Scherer, Jill A. Mikucki, and Ross D. Powell. Did holocene climate changes drive west antarctic grounding line retreat and readvance? The Cryosphere, 15(10):4655–4673, October 2021.

[66] M. E. Weber, P. U. Clark, G. Kuhn, A. Timmermann, D. Sprenk, R. Gladstone, X. Zhang, G. Lohmann, L. Menviel, M. O. Chikamoto, T. Friedrich, and C. Ohlwein. Millennial-scale variability in antarctic ice-sheet discharge during the last deglaciation. Nature, 510(7503):134–138, May 2014.

[67] Anna-Mireilla Hayden and Christine F. Dow. Examining the effect of ice dynamic changes on subglacial hydrology through modelling of a synthetic antarctic glacier. Journal of Glaciology, 69(278):1846–1859, September 2023.

[68] J. Jouzel, V. Masson-Delmotte, O. Cattani, G. Dreyfus, S. Falourd, G. Hoffmann, B. Minster, J. Nouet, J. M. Barnola, J. Chappellaz, H. Fischer, J. C. Gallet, S. Johnsen, M. Leuenberger, L. Loulergue, D. Luethi, H. Oerter, F. Parrenin, G. Raisbeck, D. Raynaud, A. Schilt, J. Schwander, E. Selmo, R. Souchez, R. Spahni, B. Stauffer, J. P. Steffensen, B. Stenni, T. F. Stocker, J. L. Tison, M. Werner, and E. W. Wolff. Orbital and millennial Antarctic climate variability over the past 800, 000 years. Science, 317(5839):793–796, August 2007.

[69] Devin N. Castendyk, Maciej K. Obryk, Sasha Z. Leidman, Michael Gooseff, and Ian Hawes. Lake Vanda: A sentinel for climate change in the McMurdo sound region of Antarctica. Global and Planetary Change, 144:213–227, September 2016.

[70] Sarah Seabrook, Cliff S. Law, Andrew R. Thurber, Yoann Ladroit, Vonda Cummings, Leigh Tait, Alicia Maurice, and Ian Hawes. Antarctic seep emergence and discovery in the shallow coastal environment. Nature Communications, 16(1), October 2025.

[71] Alexander B. Michaud, John E. Dore, Amanda M. Achberger, Brent C. Christner, Andrew C. Mitchell, Mark L. Skidmore, Trista J. Vick-Majors, and John C. Priscu. Microbial oxidation as a methane sink beneath the West Antarctic Ice Sheet. Nature Geoscience, 10(8):582–586, July 2017.

[72] Bruce M. Jakosky, Noora R. Alsaeed, Eryn M. Cangi, Michael S. Chaffin, Justin Deighan, Margaret E. Landis, Michael T. Mellon, and Edward M.B. Thiemann. The history of Martian water during the Hesperian and Amazonian epochs. Icarus, 443:116782, January 2026.

[73] Jeffrey S. Kargel and Robert G. Strom. Ancient glaciation on Mars. Geology, 20(1):3, 1992.

[74] John W. Holt, Ali Safaeinili, Jeffrey J. Plaut, James W. Head, Roger J. Phillips, Roberto Seu, Scott D. Kempf, Prateek Choudhary, Duncan A. Young, Nathaniel E. Putzig, Daniela Biccari, and Yonggyu Gim. Radar sounding evidence for buried glaciers in the southern mid-latitudes of Mars. Science, 322(5905):1235–1238, November 2008.

[75] Sebastian Emanuel Lauro, Elena Pettinelli, Graziella Caprarelli, Luca Guallini, Angelo Pio Rossi, Elisabetta Mattei, Barbara Cosciotti, Andrea Cicchetti, Francesco Soldovieri, Marco Cartacci, Federico Di Paolo, Raffaella Noschese, and Roberto Orosei. Multiple subglacial water bodies below the south pole of Mars unveiled by new MARSIS data. Nature Astronomy, 5(1):63–70, September 2020.

[76] Lu Pan, John Carter, Cathy Quantin-Nataf, Maxime Pineau, Boris Chauvire, Nicolas Mangold, Laetitia Le Deit, Benjamin Rondeau, and Vincent Chevrier. Voluminous silica precipitated from Martian waters during late-stage aqueous alteration. The Planetary Science Journal, 2(2):65, April 2021.

[77] A. M. Rutledge, B. H. N. Horgan, J. R. Havig, E. B. Rampe, N. A. Scudder, and T. L. Hamilton. Silica dissolution and precipitation in glaciated volcanic environments and implications for mars. Geophysical Research Letters, 45(15):7371–7381, August 2018.

[78] Gunter Faure, Jochen Hoefs, Lois M. Jones, John B. Curtis, and Douglas E. Pride. Extreme 18o depletion in calcite and chert clasts from the Elephant Moraine on the East Antarctic ice sheet. Nature, 332(6162):352–354, March 1988.

[79] Yetang Wang, Shugui Hou, Valerie Masson-Delmotte, and Jean Jouzel. A new spatial distribution map of delta18O in Antarctic surface snow. Geophysical Research Letters, 36(6), March 2009.

[80] Martin Werner, Jean Jouzel, Valerie Masson-Delmotte, and Gerrit Lohmann. Reconciling glacial Antarctic water stable isotopes with ice sheet topography and the isotopic paleothermometer. Nature Communications, 9(1), August 2018.

[81] Gideon M. Henderson, Brenda L. Hall, Andrew Smith, and Laura F. Robinson. Control on (234U/238U) in lake water: A study in the Dry valleys of Antarctica. Chemical Geology, 226(3–4):298–308, February 2006.

[82] W. Berry Lyons, Jill A. Mikucki, Laura A. German, Kathleen A. Welch, Susan A. Welch, Christopher B. Gardner, Slawek M. Tulaczyk, Erin C. Pettit, Julia Kowalski, and Bernd Dachwald. The geochemistry of englacial brine from Taylor Glacier, Antarctica. Journal of Geophysical Research: Biogeosciences, 124(3):633–648, March 2019.

[83] Peter M. Chutcharavan, Andrea Dutton, and Michael J. Ellwood. Seawater 234u/238u recorded by modern and fossil corals. Geochimica et Cosmochimica Acta, 224:1–17, March 2018.

[84] Denis Lacelle. Environmental setting, (micro)morphologies and stable c–o isotope composition of cold climate carbonate precipitates—a review and evaluation of their potential as paleoclimatic proxies. Quaternary Science Reviews, 26(11–12):1670–1689, June 2007.

[85] Berry Lyons, Kelly Foley, Anne Carey, Melisa Diaz, Gabriel Bowen, and Thure Cerling. The isotopic geochemistry of CaCO3 encrustations in Taylor Valley, Antarctica: Implications for their origin. Acta geographica Slovenica, 60(2):125–139, December 2020.

[86] Nadin Rohland, Isabelle Glocke, Ayinuer Aximu-Petri, and Matthias Meyer. Extraction of highly degraded DNA from ancient bones, teeth and sediments for high-throughput sequencing. Nature Protocols, 13(11):2447–2461, October 2018.

[87] Joshua D Kapp, Richard E Green, and Beth Shapiro. A fast and efficient single-stranded genomic library preparation method optimized for ancient DNA. Journal of Heredity, 112(3):241–249, March 2021.

[88] Shifu Chen. Ultrafast one-pass fastq data preprocessing, quality control, and deduplication using fastp. iMeta, 2(2), May 2023.

[89] Jared T. Simpson and Richard Durbin. Efficient de novo assembly of large genomes using compressed data structures. Genome Research, 22(3):549–556, December 2011.

[90] S Andrews. Fastqc: A quality control tool for high throughput sequence data. Available online at: http://www.bioinformatics.babraham.ac.uk/projects/fastqc/, 2010.

[91] Ben Langmead and Steven L Salzberg. Fast gapped-read alignment with bowtie 2. Nature Methods, 9(4):357–359, March 2012.

[92] Eric W Sayers, Evan E Bolton, J Rodney Brister, Kathi Canese, Jessica Chan, Donald C Comeau, Ryan Connor, Kathryn Funk, Chris Kelly, Sunghwan Kim, Tom Madej, Aron Marchler-Bauer, Christopher Lanczycki, Stacy Lathrop, Zhiyong Lu, Francoise Thibaud-Nissen, Terence Murphy, Lon Phan, Yuri Skripchenko, Tony Tse, Jiyao Wang, Rebecca Williams, Barton W Trawick, Kim D Pruitt, and Stephen T Sherry. Database resources of the national center for biotechnology information. Nucleic Acids Research, 50(D1):D20–D26, December 2021.

[93] Petr Danecek, James K Bonfield, Jennifer Liddle, John Marshall, Valeriu Ohan, Martin O Pollard, Andrew Whitwham, Thomas Keane, Shane A McCarthy, Robert M Davies, and Heng Li. Twelve years of SAMtools and BCFtools. GigaScience, 10(2), 02 2021. giab008.

[94] Nick Youngblut, Maxime Borry, Daniel Portik, and Wei Shen. nick-youngblut/gtdb to taxdump: Improved docs, 2023.

[95] Yucheng Wang, Thorfinn Sand Korneliussen, Luke E. Holman, Andrea Manica, and Mikkel Winther Pedersen. ngsLCA — A toolkit for fast and flexible lowest common ancestor inference and taxonomic profiling of metagenomic data. Methods in Ecology and Evolution, 13(12):2699–2708, October 2022.

[96] Antonio Fernandez-Guerra, Lars Wormer, Guillaume Borrel, Tom O Delmont, Bo Elberling, Marcus Elvert, A. Murat Eren, Simonetta Gribaldo, Rasmus Amund Henriksen, Kai-Uwe Hinrichs, Annika Jochheim, Thorfinn S. Korneliussen, Mart Krupovic, Nicolaj K. Larsen, Rafael Perez-Laso, Mikkel Winther Pedersen, Vivi K. Pedersen, Anthony H. Ruter, Karina K. Sand, Martin Sikora, Martin Steinegger, Iva Veseli, Yucheng Wang, Lei Zhao, Marina Zure, Kurt H. Kjær, and Eske Willerslev.̌ Two-million-year-old microbial communities from the Kap København formation in North Greenland. June 2023.

[97] Yi Wang, David Schleheck, Elena Marinova, Martin Wessels, Sebastian Schaller, Flavio S. Anselmetti, Antje Schwalb, Mikkel W. Pedersen, and Laura S. Epp. Microbial sedimentary DNA from a cultural landscape disentangles the impacts of humans and nature over the past 13.5 thousand years. March 2025.

[98] Bianca De Sanctis, Cade Mirchandi, Haoran Dong, Ruairidh MacLeod, Russ Corbett-Detig, and Yucheng Wang. Bamdam: A post-mapping authentication toolkit for ancient metagenomics. Genome Biology, In Press. 2025. GitHub: https://github.com/bdesanctis/bamdam/.

[99] Raphael Eisenhofer, Jeremiah J. Minich, Clarisse Marotz, Alan Cooper, Rob Knight, and Laura S. Weyrich. Contamination in low microbial biomass microbiome studies: Issues and recommendations. Trends in Microbiology, 27(2):105–117, February 2019.

[100] Nicole S. Treichel, Thomas C. A. Hitch, John Penders, and Thomas Clavel. Benchmarking of shotgun sequencing depth highlights strain-level limitations of metagenomic analysis. March 2025.

[101] Jim Shaw and Yun William Yu. Rapid species-level metagenome profiling and containment estimation with sylph. Nature Biotechnology, October 2024.

[102] Jari Oksanen, Gavin L. Simpson, F. Guillaume Blanchet, Roeland Kindt, Pierre Legendre, Peter R. Minchin, R.B. O’Hara, Peter Solymos, M. Henry H. Stevens, Eduard Szoecs, Helene Wagner, Matt Barbour, Michael Bedward, Ben Bolker, Daniel Borcard, Tuomas Borman, Gustavo Carvalho, Michael Chirico, Miquel De Caceres, Sebastien Durand, Heloisa Beatriz Antoniazi Evangelista, Rich FitzJohn, Michael Friendly, Brendan Furneaux, Geoffrey Hannigan, Mark O. Hill, Leo Lahti, Dan McGlinn, Marie-Helene Ouellette, Eduardo Ribeiro Cunha, Tyler Smith, Adrian Stier, Cajo J.F. Ter Braak, and James Weedon. vegan: Community Ecology Package, 2025. R package version 2.7-0, https://github.com/vegandevs/vegan.

[103] Isabel Schober, Julia Koblitz, Joaquim Sarda Carbasse, Christian Ebeling, Marvin Leon Schmidt, Adam Podstawka, Rohit Gupta, Vinodh Ilangovan, Javad Chamanara, Jorg Overmann, and Lorenz Christian Reimer. Bacdive in 2025: the core database for prokaryotic strain data. Nucleic Acids Research, 53(D1):D748–D756, October 2024.

[104] Hadley Wickham, Mara Averick, Jennifer Bryan, Winston Chang, Lucy D’Agostino McGowan, Romain Francois, Garrett Grolemund, Alex Hayes, Lionel Henry, Jim Hester, Max Kuhn, Thomas Lin Pedersen, Evan Miller, Stephan Milton Bache, Kirill Muller, Jeroen Ooms, David Robinson, Dana Paige Seidel, Vitalie Spinu, Kohske Takahashi, Davis Vaughan, Claus Wilke, Kara Woo, and Hiroaki Yutani. Welcome to the tidyverse. Journal of Open Source Software, 4(43):1686, 2019.

[105] Philippe Massicotte and Andy South. rnaturalearth: World Map Data from Natural Earth, 2023. R package version 1.0.1.

[106] Edzer Pebesma. Simple Features for R: Standardized Support for Spatial Vector Data. The R Journal, 10(1):439–446, 2018.

