## Supplementary Information for "Ancient metagenomics reveals subglacial microbiomes driven by oxygen availability"

#### 1 Subglacial precipitate subsampling and sequencing details

We sequenced 25 subsamples (referred to as 'samples' throughout) from 22 total subglacial precipitates, plus a modern Antarctic surface control rock, across 3 separate single-stranded sequencing rounds. We also sequenced 2-3 negative controls (extraction blanks and library controls) per sequencing round, yielding 7 negative control libraries altogether. This yielded a total of 35 sequencing libraries. In most cases, we sampled a single rock and created a single sequencing library, but some were duplicated within or across rounds, for two reasons. First, almost all of the libraries were created after bathing the rocks in bleach, except for S2 (from MA113), S31 (from 19ACAG81), S5 (from MV2), and S7 (from PRR52588). In each of these cases, we separately processed separate bleached and unbleached libraries through our entire bioinformatics mapping pipeline. We found that bleaching samples made essentially no difference in either community composition or damage. Because of this, we combined these libraries with their matched bleach-treated libraries. Second, a few rocks were subsampled twice, from different sections. These libraries sometimes yielded sufficiently different results that we decided to keep these different subsamples from the same rock separate through the entire pipeline, and not merge them. We sampled both opal and calcite sections from the rock MA113, yielding the samples MA1 and MA2, and we sequenced from PRR50504 a bulk calcite sample (EM2), an organic calcite sample from the top (EM3), and a second calcite sample from the top (EM4), all of which we kept separate in the end. As can be seen in Supplementary Figure 1, these rock sections differ visually, and their microbial communities may differ as well (see Krona plots). Further details can be found in Supplementary Data 1.1.

#### 2 Bioinformatic decontamination and taxonomic assignment

In order to stringently decontaminate our samples, we extracted all 930 taxonomic nodes in the GTDB taxonomy with at least 50 aggregated reads over 7 negative controls. For context, the most abundant negative control genera (with more than 1000 assigned reads in at least one negative control sample) were the human or skin associated bacteria *Cutibacterium*, *Staphylococcus*, and *Corynebacterium*, lab contaminants *Bradyrhizobium*, *Sphingomonas*, *Acinetobacter*, *Leuconostoc*, *Xanthomonas*, and *Pseudomonas*, and lastly *Calothrix*, a freshwater cyanobacteria. For the most part these are well-known contaminants (1). Reads which were assigned to any of these 930 taxonomic nodes were then removed in the samples in bamdam shrink, and then reaggregated up the taxonomy. This means that, if two different species of the same genus were present in both a negative control and a sample, we would remove all reads in the sample mapping to both the genus and the the negative control species, but not to the species present in only the sample. The reads for that species would then count towards the genus-level read counts in the sample. In this way, we keep all reads mapping to sufficiently specific taxonomic nodes that do not appear in the negative controls. For example, we end up keeping a species of *Corynebacterium* which was in the samples but not identified in the negative controls – after all, the samples and surface control were handled by more humans and exposed to more human-associated environments than the negative controls, and they were sequenced deeper – which is why we see the *Corynebacterium* genus on the far left of Figure 3. But other than these rare cases, this is actually the behavior we want, since for example, there are certainly subglacial taxa and negative control taxa within the same phylum. Supplementary Figure 2 shows the results of this bioinformatic decontamination

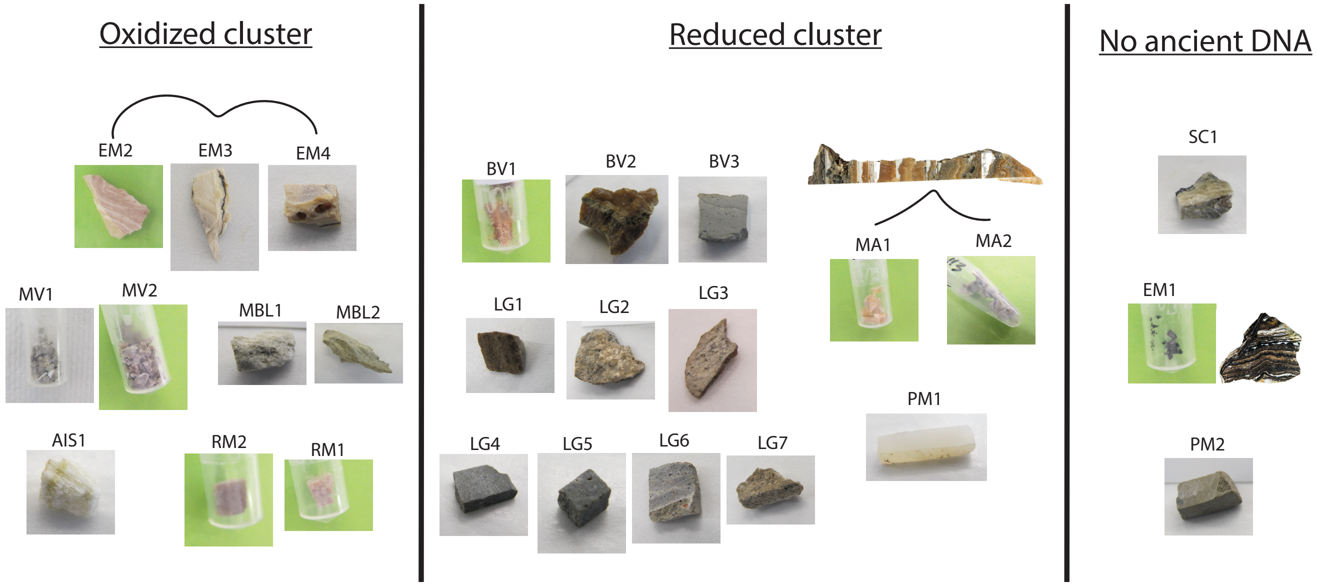


**Figure 1:** Laboratory pictures of each rock sample, organized by our designation of oxidized (left), reduced (center) or containing no detectable ancient DNA (right). Samples from different sections of the same subglacial precipitate are connected.

procedure. We keep the vast majority of taxa, and usually keep the majority of reads, although in some cases (AIS1, EM1-4, PM2) this is a minority, indicating a low amount of endogenous DNA and/or a high degree of laboratory contamination. EM1 and PM2 had lowest number of reads after filtering, and unsurprisingly did not contain any ancient DNA. For EM1, this may be because of low sample availability, as can be seen in Supplementary Figure 1.

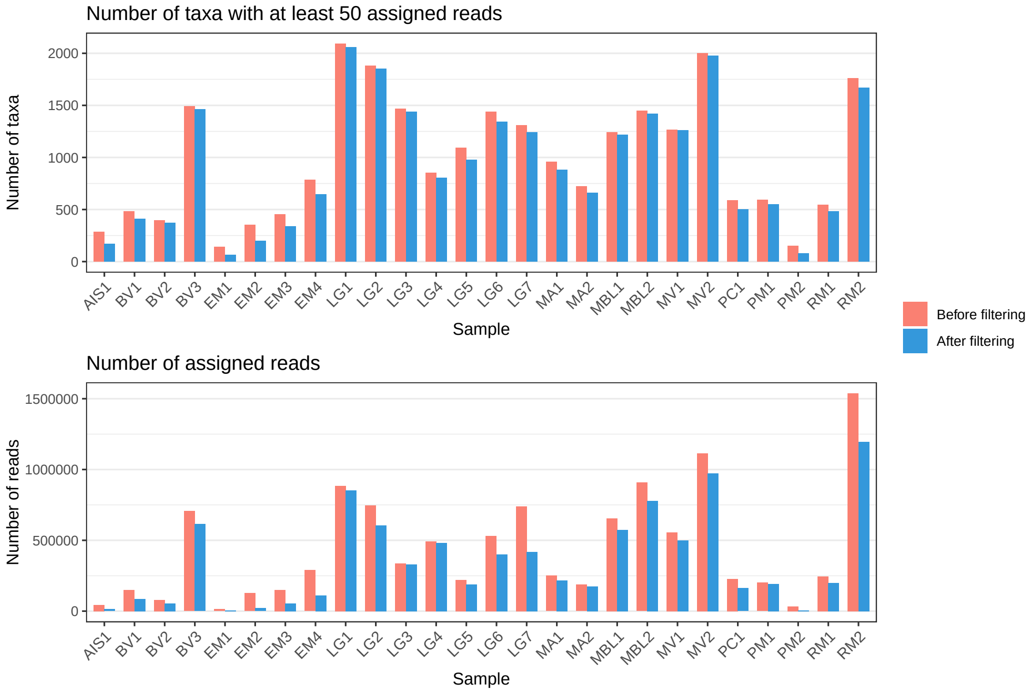


**Figure 2:** Effect of bioinformatic decontamination, removing sample reads which are assigned to taxa prevalent in the negative controls.

### 3 An extended analysis of taxa by damage

Here we present a series of median damage by taxa plots such as in main Figure 3A, but across different taxonomic levels and read count thresholds. In all cases, the surface control is shown as red dots. First, Supplementary Figure 3 shows an analog of main Figure 3 but at class level. The 19 abundant classes in this figure to the right of the rightmost red point are considered 'subglacial classes' for the purposes clustering the subglacial communities, in main Section 4. Choosing class as a taxonomic level to define 'subglacial taxa' for the clustering section is somewhat arbitrary, as all taxonomic levels define some ancient and some modern taxa, but we found that overall, class level gave us the most total reads while still retaining a clear damage separation and ecological profile. Supplementary Figures 4 and 5 then show the effect of lowering the read threshold for genus and class level. If the reader desires further intuition for these samples and their detailed taxonomic makeup, we recommend exploring the interactive Krona plots. Relevant files are also available on the Github at `www.github.com/bdesanctis/subglacial-precipitates`.

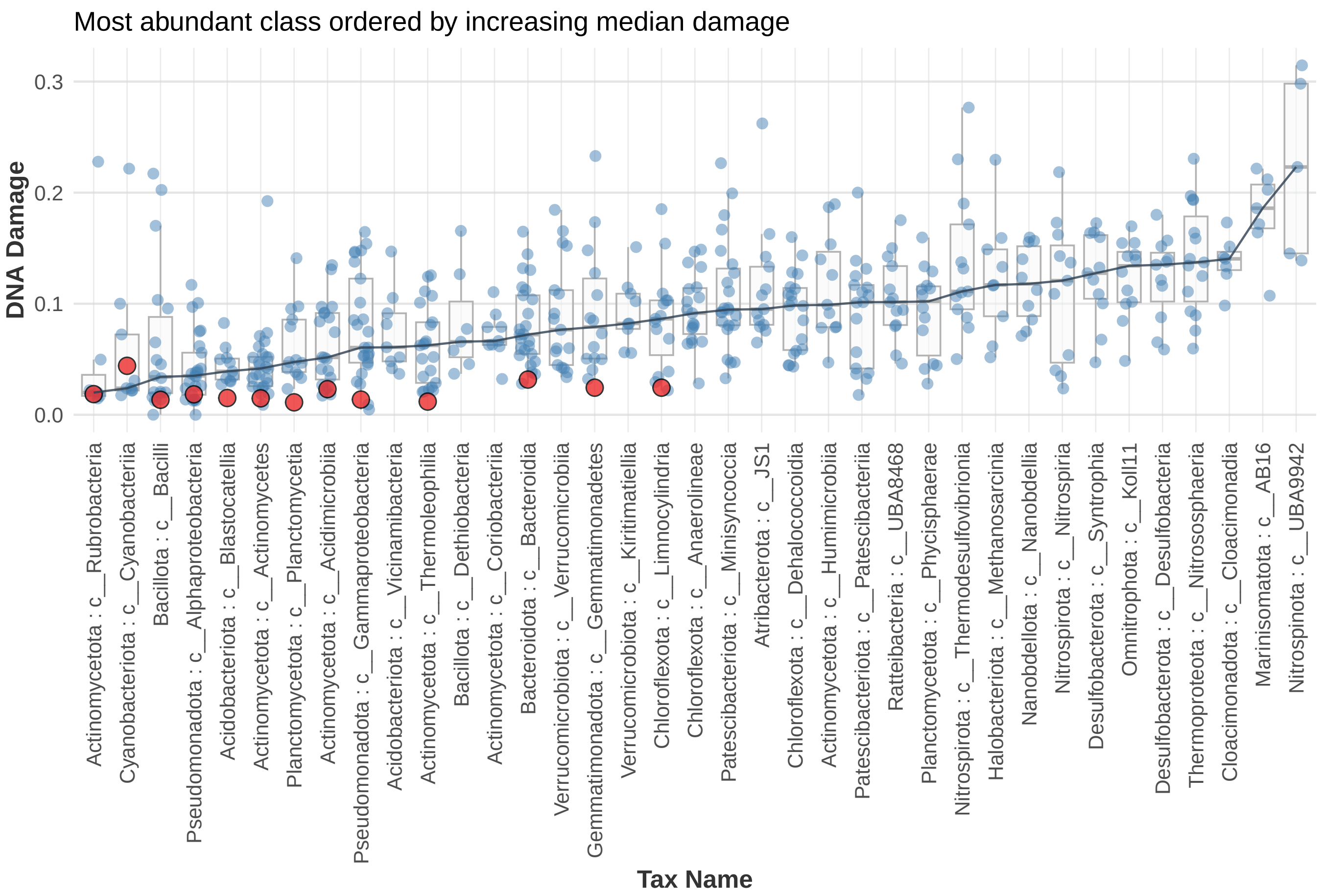


**Figure 3:** Classes with more than 30,000 reads across samples ordered by median damage. Points are only shown for, and damage is only calculated on, samples with at least 200 reads for that class. The positive control damage is not included in the median damage calculation. We only show taxa present with more than 200 reads in at least 3 samples. Blue dots are precipitate samples and red dots are the surface control.

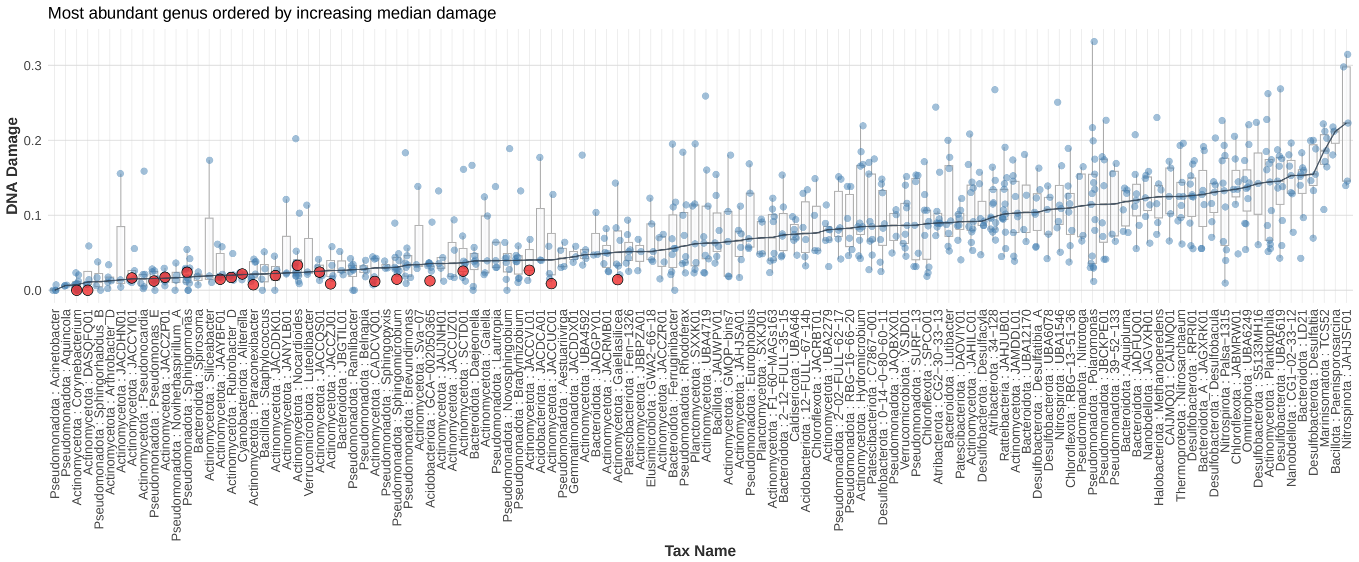


**Figure 4:** A version of main Figure 3 but with a lower total read threshold. Genera with more than 10,000 reads across samples ordered by median damage. Points are only shown for, and damage is only calculated on, samples with at least 200 reads for that genus. The positive control damage is not included in the median damage calculation. We only show taxa present with more than 200 reads in at least 3 samples. Blue dots are precipitate samples and red dots are the surface control.

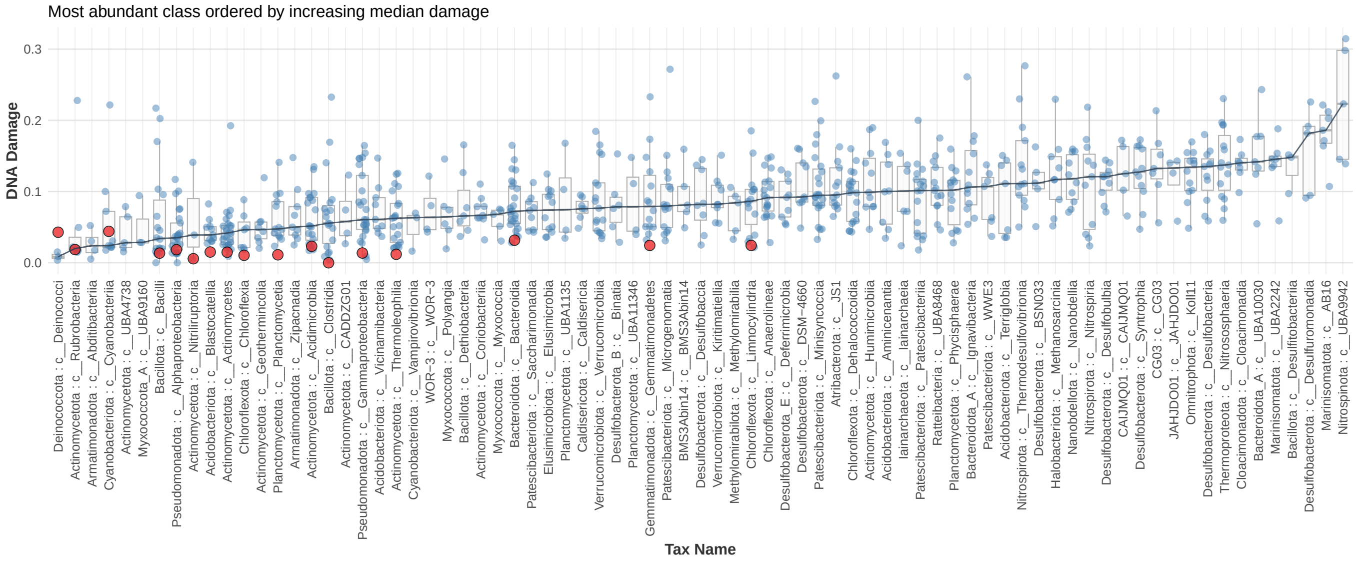


**Figure 5:** Classes with more than 10,000 reads across samples ordered by median damage. Points are only shown for, and damage is only calculated on, samples with at least 200 reads for that class. The positive control damage is not included in the median damage calculation. We only show taxa present with more than 200 reads in at least 3 samples. Blue dots are precipitate samples and red dots are the surface control.

### 4 Clustering the subglacial communities

Supplementary Figure 6 shows the loadings per taxonomic node for the subglacial NMDS in main Figure 4A. Each point is a taxonomic node. Note that the point sizes indicate the total reads assigned to that node or underneath it – that is, the total aggregated reads – but the actual NMDS was performed on the unaggregated number of reads to each taxonomic node, so as not to double-count reads. We see that, even within the subglacial classes which are shared between the clusters, there is still usually a split at more specific levels.

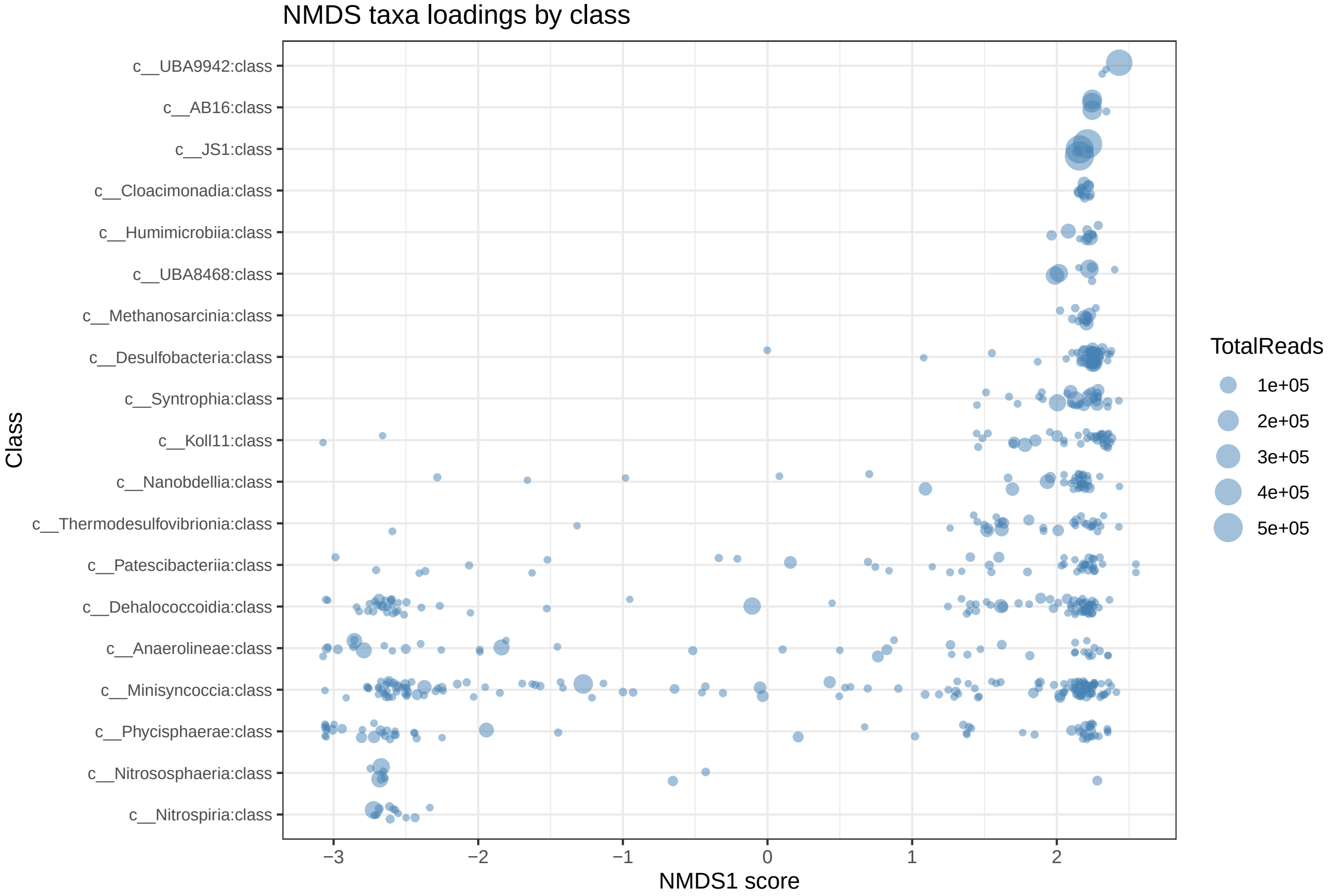


**Figure 6:** The loadings per taxonomic node in the subglacial NMDS in main Figure 4a, grouped by class.

### 5 Contextualizing previously published direct samples

#### 5.1 Subglacial Lake Whillans, Subglacial Lake Mercer, and Blood Falls Brines

Throughout this paper, we compare to three previously published, directly sampled Antarctic subglacial sites: Subglacial Lake Whillans, Subglacial Lake Mercer, and Blood Falls Brines. All microbial studies on these sites thus far did not use shotgun metagenomics as done here, so our data were not directly comparable. In particular, material from all three of these Antarctic subglacial sites has been sequenced using amplicons of the 16S rRNA gene (2; 3; 4; 5), and more recently SLM with single-cell sequencing (6). Furthermore, some of their top reported taxa are not in our subglacial classes (as expected; see the end of Section 3). Therefore, to do this comparison, we used their 'top taxa', which in many cases did not match the nomenclature within the GTDB 226.0 taxonomy used in this paper. Here, we detail exactly how we reconciled names for the top taxa from each of these sites, to create taxa lists which were usable for this paper.

Subglacial Lake Mercer was the most straightforward of the three, because there was a single-cell sequencing paper published recently on which we could rely and which used a recent version of the GTDB taxonomy (v214) (6). We used the taxa reported in Figure 1 of this paper, that is, the highly abundant genera. Conveniently, all names translate directly to the GTDB taxonomy version used here (v226.0). The original paper also reports several families and higher taxonomic levels, but we chose to restrict ourselves to genus level. There has also been 16S metabarcoding work done on SLM previously, but with less taxonomic resolution than this single-cell study, and not based on the GTDB taxonomy (5), so we relied on the (6) study for comparison with SLM.

Subglacial Lake Whillans was analyzed in a previous study with 16S metabarcoding. The top taxa we used can be seen in Fig 2 of (2). For the most part, it was straightforward to match genus names to the GTDB taxonomy we used. However, we could not find matches for *Feriphaselus* or *Thermomarinilacea*, and therefore excluded them. We also moved IDs up to the genus level, since many species listed in the original study do not have exact analogs in GTDB. We used *g__Rhodoferax* for *Albidiferax*, because in GTDB we saw that *s__Albidiferax* was in the *g__Rhodoferax* genus. Once again, we restricted ourselves to genus level.

Blood Falls Brine was slightly more complex. We relied on the 16S taxonomic identifications from Figure 4 in (4), in particular those taxa found in the EMB (end member brine) community, indicated in the original figure by the green EMB column. For *DHVE6_ge*, the original study said "The SILVA (v128) reference taxonomy contains sequences from the *DHVE6* archaeal group classified as *Woesearchaeota* but also containing sequences closer to *Pacearchaeota*." We found little information on *DHVE6_ge* otherwise, and so we used both *o__Pacearchaeales* and *o__Woesarcheaeles* to replace it. For *NB1-n_ge*, we relied on (7), which suggested the new name *Candidatus 'Izimaplasma'* for this clade, and we accordingly used *f__Izemoplasmataceae* in the GTDB taxonomy. We mapped *Sva0996_marine_group_ge* to *'s__marine group bacterium Sva0996 bin134'* in GTDB, then bumped it up to its genus level, which was *g__Poriferisocius*. There was no GTDB result for *Maritimimonas*, but we found an LPSN page `https://lpsn.dsmz.de/genus/maritimimonas` which states the parent taxon of *Maritimimonas* as *Flavobacteriaceae*, and so we used the GTDB taxon *o__Flavobacteriales*, just to be safe. We did not include uninformative identifications from the original paper, namely *Bacteria_unc*, *Deltaproteobacteria_unc*, and *Bacteroidetes_unc*.

Finally, we discuss the checkmarks in our Main Figure 1. Most of the relevant taxa were already mentioned above. Furthermore, we put a checkmark for BFB under our *Atribacterota* genus *34-128* because the BFB 16S paper (4) reported the same *Atribacterota* class *c__JS1* (in the text). We also put a check mark on our genus *g__Desulfobacula* for BFB because it is inside the family *f__Desulfobacteraceae*, which was identified as an abundant taxa in Blood Falls Brine. Lastly, *Lutibacter* was found in a previous SLM 16S paper (5).

A side effect of placing SLM, SLW and BFB into our existing clusters is the identification of potential abundant lab or surface contaminants in these studies. Main Figure 4D has a column for SC1, the surface control. We see that some taxa appear with nontrivial abundance in the surface control: g__F1-60-MAGs163 from SLM, g__Methylobacter and g__Acinetobacter (and, less so, g__Dietzia) from SLW, and o__Flavobacteriales and f__Xanthomonadacea from BFB. This hypothesis is further supported by low damage for these taxa in our samples (e.g. see Krona plots), and the fact that removing these potential contaminant taxa would clean up our cluster placement. Of particular interest in this list is g__Methylobacter. There are Methylobacter-associated genomes in both SLM and SLW (though it was not a 'top taxa' in SLM, so is not shown in Main Figure 4E), with genes associated to methane oxidation in the former (6; 2). Although we do find ~3500 Methylobacter reads across samples, they are almost certainly contamination, as they are undamaged (mean 0.01 C-to-T frequency), present in the surface control, and even present in our negative controls - notably, Methylobacter has been identified as a contaminant of DNA extraction kit reagents (8) and has previously been associated with the MQ ultrapure water from Crary Lab (i.e., water collected directly from the MQ ultrapure water system at McMurdo Station, Antarctica (9). All 3 sampling projects (SLM, SLW and BFB) originated out of McMurdo Station. Another possibility is that these taxa are genuinely present both in our controls and in SLM, SLW or BFB.

Lastly, we predicted oxygen tolerance for each of these SLM, SLW and BFB associated taxa in the GTDB 226.0 taxonomy, shown in the tables below. To do this, we collected all genomes from GTDB (10) with associated protein sequences underneath these taxonomic nodes, ran GenomeSPOT (11), and extracted the oxygen tolerance (0 for anaerobic or 1 for aerobic) per genome. Then to estimate oxygen tolerance for each taxonomic node in the tables below, we averaged those values estimated on the genomes underneath them. Oxygen tolerance is more frequently predicted in SLW and SLM taxa than BFB, further reinforcing the cluster memberships we find in Figure 4.

| Source | Original name | Name used in GTDB 226.0 | Citation | Oxygen tolerance |
|---|---|---|---|---|
| SLM | g__SYFI01 | g__SYFI01 | Kim et al 2025 | 0.4 |
| SLM | g__F1-60-MAGs163 | g__F1-60-MAGs163 | Kim et al 2025 | 1 |
| SLM | g__F1-60-MAGs149 | g__F1-60-MAGs149 | Kim et al 2025 | 0.71 |
| SLM | g__UBA4592 | g__UBA4592 | Kim et al 2025 | 1 |
| SLM | g__UBA10799 | g__UBA10799 | Kim et al 2025 | 1 |
| SLM | g__Planktophila | g__Planktophila | Kim et al 2025 | 1 |
| SLM | g__UBA3006 | g__UBA3006 | Kim et al 2025 | 1 |
| SLM | g__RBG-16-66-20 | g__RBG-16-66-20 | Kim et al 2025 | 0.74 |
| SLM | g__PALSA-1004 | g__PALSA-1004 | Kim et al 2025 | 0.67 |
| SLM | g__Polaromonas | g__Polaromonas | Kim et al 2025 | 1 |
| SLM | g__Nitrotoga | g__Nitrotoga | Kim et al 2025 | 1 |
| SLM | g__39-52-133 | g__39-52-133 | Kim et al 2025 | 1 |
| SLM | g__SURF-13 | g__SURF-13 | Kim et al 2025 | 1 |
| SLM | g__12-FULL-67-14b | g__12-FULL-67-14b | Kim et al 2025 | 1 |
| SLM | g__SPCO01 | g__SPCO01 | Kim et al 2025 | 1 |
| SLM | g__C7867-001 | g__C7867-001 | Kim et al 2025 | 1 |
| SLM | g__UBA1550 | g__UBA1550 | Kim et al 2025 | 0.95 |
| SLM | g__Nitrosarchaeum | g__Nitrosarchaeum | Kim et al 2025 | 0 |

**Table 1:** Name translations for taxa found in Subglacial Lake Mercer into the GTDB v226.0 taxonomy.

| Source | Original name | Name used in GTDB 226.0 | Citation | Oxygen tolerance |
|---|---|---|---|---|
| SLW | Sideroxydans lithotrophicus | g__Sideroxyarcus | Achberger et al 2016 | 1 |
| SLW | Feriphaselus amnicola | NA | Achberger et al 2016 | NA |
| SLW | Albidiferax ferrireducens | g__Rhodoferax | Achberger et al 2016 | 1 |
| SLW | Thiobacillus denitrificans | g__Thiobacillus | Achberger et al 2016 | 1 |
| SLW | Acidiferrobacter thiooxydans | g__Acidiferrobacter | Achberger et al 2016 | 1 |
| SLW | Candidatus Nitrotoga arctica | g__Nitrotoga | Achberger et al 2016 | 1 |
| SLW | Jettenia asiatica | g__Jettenia | Achberger et al 2016 | 0 |
| SLW | Nitrosoarchaeum koreensis | g__Nitrosarchaeum | Achberger et al 2016 | 0 |
| SLW | Nitrosospira multiformis | g__Nitrosospira | Achberger et al 2016 | 1 |
| SLW | Methylobacter tundripaludum | g__Methylobacter | Achberger et al 2016 | 1 |
| SLW | Methylobacillus glycogenes | g__Methylobacillus | Achberger et al 2016 | 1 |
| SLW | Methyloversatilis thermotolerans | g__Methyloversatilis | Achberger et al 2016 | 1 |
| SLW | Polaromonas glacialis | g__Polaromonas | Achberger et al 2016 | 1 |
| SLW | Aggregicoccus edonensis | g__Aggregicoccus | Achberger et al 2016 | 1 |
| SLW | Solitalea koreensis | g__Solitalea | Achberger et al 2016 | 1 |
| SLW | Ohtaekwangia koreensis | g__Ohtaekwangia | Achberger et al 2016 | 1 |
| SLW | Acinetobacter lwoffii | g__Acinetobacter | Achberger et al 2016 | 1 |
| SLW | Smithella propionica | g__Smithella | Achberger et al 2016 | 0 |
| SLW | Candidatus Planktophila limnetica | g__Planktophila | Achberger et al 2016 | 1 |
| SLW | Ignavibacterium album | g__Ignavibacterium | Achberger et al 2016 | NA |
| SLW | Ilumatobacter fluminis | g__Ilumatobacter | Achberger et al 2016 | 1 |
| SLW | Dietzia alimentaria | g__Dietzia | Achberger et al 2016 | 1 |
| SLW | Thermomarininilacea lacunofontalis | NA | Achberger et al 2016 | NA |

**Table 2:** Name translations for taxa found in Subglacial Lake Whillans into the GTDB v226.0 taxonomy.

| Source | Original name | Name used in GTDB 226.0 | Citation | Oxygen tolerance |
|---|---|---|---|---|
| BFB | Xanthomonadaceae_unc | f__Xanthomonadaceae | Campen et al 2019 | 1 |
| BFB | Thiomicrospira | f__Thiomicrospiraceae | Campen et al 2019 | 1 |
| BFB | Flavobacterium | o__Flavobacteriales | Campen et al 2019 | 0.99 |
| BFB | NB1-n_ge | f__Izemoplasmataceae | Campen et al 2019 | 0.33 |
| BFB | Bacteria_unc | NA | Campen et al 2019 | NA |
| BFB | DHVE6_ge | o__Woesearchaeales | Campen et al 2019 | 0.02 |
| BFB | DHVE6_ge | o__Pacearchaeales | Campen et al 2019 | 0 |
| BFB | Lutibacter | g__Lutibacter | Campen et al 2019 | 1 |
| BFB | Deltaproteobacteria_unc | NA | Campen et al 2019 | NA |
| BFB | Desulfobacteraceae_unc | f__Desulfobacteraceae | Campen et al 2019 | 0 |
| BFB | Geopsychrobacter | f__Geopsychrobacteraceae | Campen et al 2019 | 0 |
| BFB | Desulfobulbaceae_unc | f__Desulfocapsaceae | Campen et al 2019 | 0 |
| BFB | Bacteroidetes_unc | NA | Campen et al 2019 | NA |
| BFB | Sva0996_marine_group_ge | g__Poriferisocius | Campen et al 2019 | 1 |
| BFB | Maritimimonas | o__Flavobacteriales | Campen et al 2019 | 0.99 |
| BFB | Atribacteria_ge | p__Atribacterota | Campen et al 2019 | 0 |

**Table 3:** Name translations for taxa found in Blood Falls Brine into the GTDB v226.0 taxonomy.

#### 5.2 A brief discussion of a previous study of subglacial precipitates, Frisia et al. 2017

Subglacial precipitates have been sequenced once previously (12). Here, the authors sequenced two 17-26 thousand year old Antarctic subglacial precipitates from Boggs Valley using 16S metabarcoding. Metabarcoding is generally considered inappropriate for ancient microbiome reconstruction, and makes it impossible to authenticate ancient DNA by assessing postmortem damage values across taxa (13). In the Frisia et al 2017 study, the authors concluded that all of their identified taxa were ancient and subglacial, and did not consider the possibility of surface contamination or compare their taxonomic results to the known Antarctic surface microbiome. Three precipitates from the Boggs Valley site were also included in our study, facilitating a comparison. In (12), the most abundant taxa in both rocks (30-40% of the precipitate microbiome) was identified as the bacterial order Thermomicrobia, which corresponds to Thermomicrobiales in the GTDB v226 taxonomy, within the class Chloroflexi. In our study, we also see more than 10,000 total reads mapping to Thermomicrobia across 8 subglacial samples, including those from both the Arctic and Antarctic, and in particular including a precipitate from the same Antarctic region, Boggs Valley. However, we find consistently low damage across these samples (a mean C-to-T frequency of 0.03), and over 2500 of the reads assigned to this order are from our Antarctic surface control. Moreover, our top genome hit for the Thermomicrobiales order is a MAG from Antarctic surface soil (14), which is what would be expected for an Antarctic surface taxa. The next most abundant bacterial taxa identified in (12) are Thermoleophilia and Euzebya, or Euzebyales in the GTDB taxonomy. Thermoleophilia composes 11% of our surface control (see Krona plots), and the Thermoleophilia genus JAAYBF01 is one of the most abundant surface genera in our samples (see Figure 3B), with consistently low damage (mean 0.04). Similarly, Euzebyales has more than 5000 reads in our samples, including one from Boggs Valley, but more than half of these reads originate from the modern surface control, and again with low damage (mean C-to-T of 0.04). Additionally, Thermomicrobiales and Thermoleophilia have previously been found in high abundance in Antarctic surface endolithic microbiomes (15). We did not attempt to systematically validate further taxa from Frisia et al 2017, but we caution that the conclusions based on microbial DNA concerning Antarctic subglacial volcanism in (12) are likely not valid. This highlights the importance of using an appropriate sequencing strategy and using damage patterns for authentication in ancient environmental DNA studies.

### 6 Gene annotation

#### 6.1 De novo assembly and functional annotation

We attempted de novo assembly and functional annotation, but this was complicated by ancient DNA limitations, which impact assembly far more than alignment (1). In particular we attempted gene annotation on de novo assembled contigs and their associated proteins for each sample using both the general-purpose eggNOG-mapper, and by comparing against targeted gene databases for sulfur, methane, nitrogen, iron, and trace gase metabolism (see Supplementary Methods for details). We attached damage values to each gene, which were inherited from their DNA parent contigs, by mapping reads back against contigs. Although there was not quite enough data to draw detailed conclusions per sample, or between clusters, results are shown in Supplementary Figure 7. In particular we show the results of the targeted analysis on methane, nitrogen and sulfur genes, and draw a surface control damage distribution in a red band from the eggNOG results. This figure includes only genes that were found in at least two samples. No iron cycling genes were found. Notably absent is mcrA, a functional marker gene for methanogenesis.

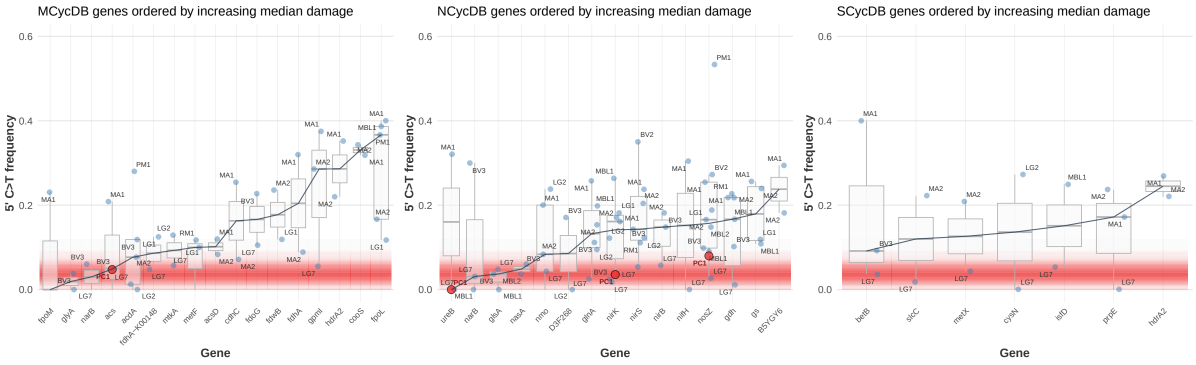


**Figure 7:** Genes related to methane, nitrogen and sulfur metabolism, ordered by median damage, in the subglacial precipitates and surface control. Genes are only shown if they are found in at least 2 samples. The horizontal red band indicates the damage distribution found from over a thousand eggNOG genes which had more than 100 recruited reads in the surface control.

### 7 Metagenome-assembled genomes

In the reduced cluster, a subset of both Arctic and Antarctic samples contains Nitrospinota, one of the most abundant nitrite oxidizers in the marine water column (16). In particular, LG4 was almost exclusively dominated by one species of Nitrospinota (over 300,000 uniquely mapped reads, e.g. see Figure 2D). Based on mapping, Nitrospinota is always found highly damaged (right of Figure 3), and in LG4 there are 332,284 reads with mean damage 0.22 mapping uniquely to the accession GCA_018830405.1, to which our MAG classifies. Incredibly, only 62 Nitrospinota reads map to anything else within the Nitrospinota phylum, which is composed of 216 representative genomes in the GTDB v226.0 reference database, indicating a high divergence from the rest of the Nitrospinota genomes. Thanks to this highly specific Nitrospinota content in LG4, we were able to reconstruct a high-quality Nitrospinota MAG (metagenome-assembled genome) from this sample, despite ancient DNA challenges typically associated with de novo assembly (1). The MAG had 89.59% completion and 0.43% contamination and was classified in the GTDB taxonomy as species JAHJSF01 sp018830405, with 97.9% ANI to reference GCA_018830405.1 (Bacteria; Nitrospinota; UBA9942; UBA9942; JAHJSF01; JAHJSF01; JAHJSF01 sp018830405), a genome isolated from deep groundwater from the Fennoscandian Shield, an isolated, oligotrophic system (17). We assessed postmortem damage on each MAG contig individually using modified bamdam code (18). Our reasoning was that, if these MAGs represent true ancient subglacial taxa, the damage should be high across all of its contigs, and at very similar values. This is indeed what we found, confirming that the MAG is ancient and subglacial. This ancient Nitrospinota MAG contained genes for near complete denitrification, including genes encoding the enzymes for the nitrate reduction, nitrite reduction, and nitric oxide reduction steps, highlighting its role in nitrogen cycling in reduced conditions. While Nitrospinota is widespread on Earth today, a recent ancestral state reconstruction study (19) determined that the last common ancestor of the Nitrospinota was most likely restricted to terrestrial subsurface aquifers, where this group relied on sulfur and hydrogen as electron sources.

Although we classified the genus Nocardioides as a surface taxa (Figure 3), there are two samples with high damage for this genus, and for the wider Nocardioidaceae family. In this same family, Nocardioides_C, a different genus to Nocardioides, is present only in samples BV1, BV2, BV3, LG2, LG7, and MBL2, with variable damage, and with a mean read-weighted damage across samples of 0.136. BV2 has 2484 reads hitting this Nocardioides_C genus and a mean of 0.22 C-to-T frequency amongst those reads. We assembled a MAG from BV2 which was classified to the UBA4001 genus of Nocardioidaceae (Bacteria;Actinomycetota; Actinomycetes; Propionibacteriales; Nocardioidaceae; Nocardioides_C). This was 97.06% complete and 3.89% contaminated. Similar to Nitrospinota above, we assessed postmortem damage on each MAG contig separately, which showed a consistent and high damage signal. We therefore tentatively suggest that this MAG is also ancient and subglacial, though we find this case less compelling than the Nitrospinota above. It is interesting that relatively few reads map to this genus, but that a MAG could be assembled that is phylogenetically assigned within it.

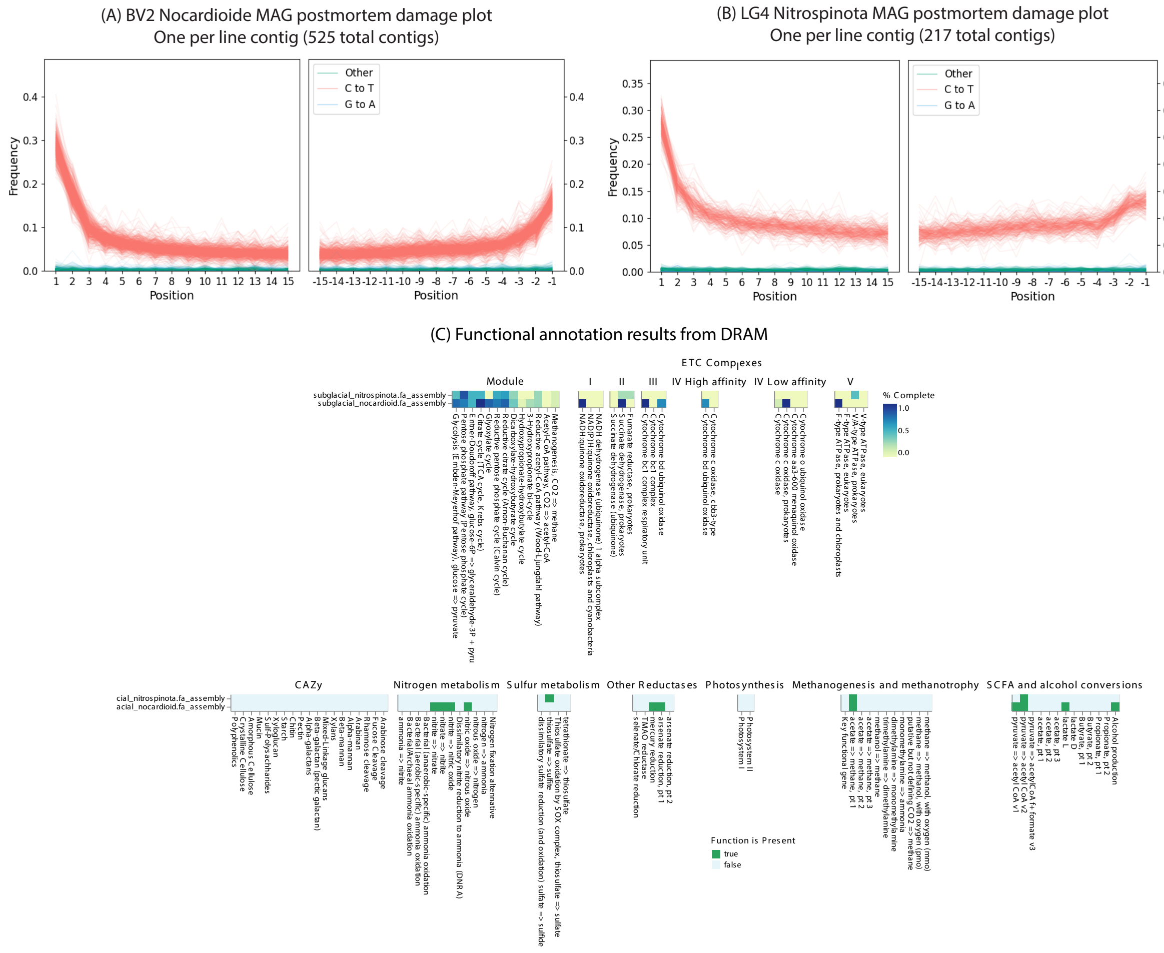


**Figure 8:** A,B: Damage plots for each assembled MAG. C: Functional annotation results from DRAM.

### 8 Precipitate formation ages

Here we show precipitate formation age versus the EDC Dome C ice core $\delta$ D, which serves as a proxy for Southern Hemisphere and Southern ocean temperature. Lower $\delta$ D values correspond to colder temperatures.

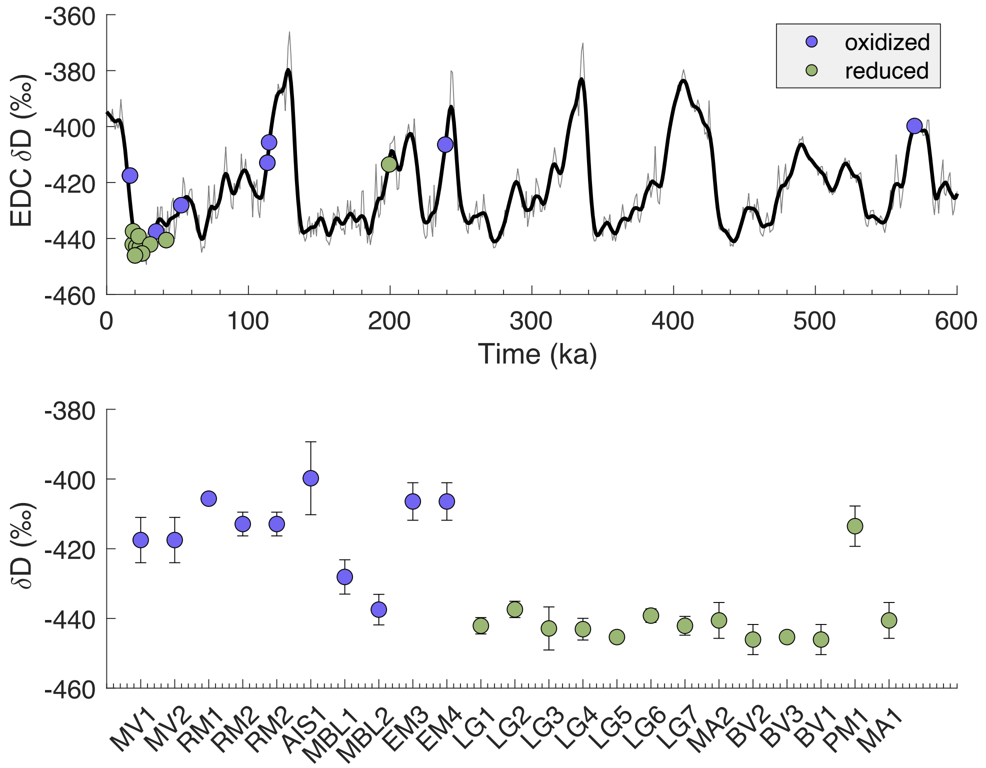


**Figure 9:** Compilation of U-Th age results for subglacial precipitates from refs (20; 21; 22; 23; 24) and this study (for sample AIS1), color coded, according to their redox grouping, as defined by taxonomic abundances, and Fe and Mn concentrations from in Figure 4 and 5 in the main text. The top panel maps the calcite precipitation ages onto the $\delta$ D (%) time series from EPICA Dome C ice core (on AICC 2012 timescale (25)). The $\delta$ D (%) value serves as proxy for Southern Hemisphere and Southern ocean temperature where isotopically light/heavy values correspond to cold/warm periods. The bottom panel provides a comparison of the $\delta$ D (%) values for the precipitate sample formation times.

### 9 Supplementary bioinformatic methods

**Assembly, gene prediction and functional annotation.** To investigate functional profiles, we ran the following pipeline on all samples plus controls, starting from the clean fastq files. We used seqtk-kit v1.4 (26) to turn clean fastqs into fasta files, then IDBA-Ud (27) with default parameters to create initial contigs. We removed all contigs under 500bp. In 3 out of 5 negative controls and 3 samples, no contigs met this filtering threshold. On the remaining controls and samples we ran Prodigal V2.6.3 (28) on metagenome mode to predict open reading frames, which yielded a CDS (coding sequence) fasta file for each sample. On these CDS files, we then ran Eggnog emapper v2.1.12 (29) with –type CDS, diamond v2.1.10 (30), and database v5.0.2, to annotate functional profiles. Clean (QCed) fastq reads were remapped against the filtered contig fastas using bwa aln v0.7.18 (31) with ancient DNA parameters (-l 1024 -n 0.01 -o 2 (32)). This was used to calculate 5' C-to-T transition frequency for each contig, by using bamdam (18) in single stranded mode and by artificially setting all the contigs to a single taxonomic level and creating a fake lca file. Each CDS region then inherited the 5' C-to-T transition frequency of its original contig. We chose to calculate damage on filtered contigs rather than CDS regions because this calculation is more accurate with more reads. We removed all genes which had less than 100 reads mapped to their parent contigs. Of the two negative controls with any eggNOG annotations, the first had only one annotation which had no KEGG pathway attached to it, and the second had 2082 annotations, of which 90% were associated with the *Corynebacteriaceae* family, a human-associated contaminant.

**Targeted gene analysis.** Targeted analysis of 762 genes related to methane cycling, sulfur cycling, nitrogen cycling, iron cycling, and trace gas metabolism were performed on the trimmed and quality filtered metagenomes. In particular, we used compilations of protein sequences following methods described in (33) and (34). These compilations of protein sequences from existing databases were searched against our IDBA-UD assembled contig files using the blastx function of DIAMOND v. 2.1.11 (30) with a query coverage of 80% (30). The reference databases used are as follows: the reference sequences of the 207 sulfur cycling genes were from SCycDB (35), the 68 N cycling genes were from NCycDB (36), the 298 methane cycling genes were from (37), iron cycling genes were from FeGenie (38). Reference databases for the acetyl-CoA-succinyl-CoA carbon fixation cycle genes (36 genes) and the C1 carbon fixation pathway genes (20 genes), described in (39), were downloaded from Uniprot (40). References sequences were restricted to reviewed entries (Swiss-Prot) unless none existed for that gene. CbbL and NifJ reference sequences were from (41). Following the methods of (34), positive hits were considered those with an identity threshold >60% and a maximum e-value threshold of $10^{-10}$. Gene hits inherited 5' C-to-T misincorporation frequencies as calculated on their parent contigs, as described above. As for the eggNOG analysis, we removed all genes which had less than 100 reads mapped to their parent contigs.

**Recovery and analysis of metagenome assembled genomes (MAGs).** Metagenome-assembled genomes (MAGs) were recovered following methods modified from (33; 42) and (43). To summarize, quality-filtered and trimmed sequences were assembled independently using three different assemblers: metaSpades v.4.0.0 (44), IDBA-UD v.1.1.3 (27), and CarpeDeam v.1.0.0 (45) with default parameters. Reads from each assembly for each sample were mapped to scaffolds using Bowtie2 v.2.5.1 (46). Abundance files were created and formatted from scaffold maps with samtools v.1.21 (47) and Anvio v.8 (48). Genome-resolved bins were obtained from scaffolds with CONCOCT v.1.1.0 (49), MaxBin2 v.2.2.4 (50), and MetaBAT2 v.2.12.1 (51). A dereplicated set of bins for each sample was then identified using DAS_Tools v1.1.7 (52), quality of the dereplicated bins was determined with CheckM v.1.2.3 (53), open reading frames were predicted with Prodigal v.2.6.3 (28)), and taxonomy was assigned to each MAG with GTDBtk v.2.4.0 using the classify workflow (54) based on the Genome Taxonomy Database (GTDB) release 02-RS220 (55). Of the MAGs assembled, just two were classified as high quality, > 90% complete and <10% contaminated (53). To determine if the two MAGs were damaged, we mapped the quality-control filtered fastqs against the assemblies from each of the two samples using bwa aln v0.7.18 (31) with ancient DNA parameters (-l 1024 -n 0.01 -o 2 (32)). As for the gene analysis above, we artifically set all the contigs of each MAG to the same taxonomic level and created a fake lca file reflecting this, then used bamdam (18) to assess damage values per contig. We slightly modified the code in bamdam plotdamage to plot lines for each contig instead of each sample. We used DRAM (56) to obtain functional annotations for each of the two MAGs.

### References


[1] Raphael Eisenhofer, Jeremiah J. Minich, Clarisse Marotz, Alan Cooper, Rob Knight, and Laura S. Weyrich. Contamination in low microbial biomass microbiome studies: Issues and recommendations. *Trends in Microbiology*, 27(2):105–117, February 2019.

[2] Amanda M. Achberger, Brent C. Christner, Alexander B. Michaud, John C. Priscu, Mark L. Skidmore, and Trista J. Vick-Majors. Microbial community structure of Subglacial Lake Whillans, West Antarctica. *Frontiers in Microbiology*, 7, September 2016.

[3] Brent C. Christner, John C. Priscu, Amanda M. Achberger, Carlo Barbante, Sasha P. Carter, Knut Christianson, Alexander B. Michaud, Jill A. Mikucki, Andrew C. Mitchell, Mark L. Skidmore, Trista J. Vick-Majors, W. P. Adkins, S. Anandakrishnan, G. Barcheck, L. Beem, A. Behar, M. Beitch, R. Bolsey, C. Branecky, R. Edwards, A. Fisher, H. A. Fricker, N. Foley, B. Guthrie, T. Hodson, H. Horgan, R. Jacobel, S. Kelley, K. D. Mankoff, E. McBryan, R. Powell, A. Purcell, D. Sampson, R. Scherer, J. Sherve, M. Siegfried, and S. Tulaczyk. A microbial ecosystem beneath the West Antarctic ice sheet. *Nature*, 512(7514):310–313, August 2014.

[4] Richard Campen, Julia Kowalski, W. Berry Lyons, Slawek Tulaczyk, Bernd Dachwald, Erin Pettit, Kathleen A. Welch, and Jill A. Mikucki. Microbial diversity of an Antarctic subglacial community and high-resolution replicate sampling inform hydrological connectivity in a polar desert. *Environmental Microbiology*, 21(7):2290–2306, April 2019.

[5] Christina L Davis, Ryan A Venturelli, Alexander B Michaud, Jon R Hawkings, Amanda M Achberger, Trista J Vick-Majors, Brad E Rosenheim, John E Dore, August Steigmeyer, Mark L Skidmore, Joel D Barker, Liane G Benning, Matthew R Siegfried, John C Priscu, Brent C Christner, Carlo Barbante, Mark Bowling, Justin Burnett, Timothy Campbell, Billy Collins, Cindy Dean, Dennis Duling, Helen A Fricker, Alan Gagnon, Christopher Gardner, Dar Gibson, Chloe Gustafson, David Harwood, Jonas Kalin, Kathy Kasic, Ok-Sun Kim, Edwin Krula, Amy Leventer, Wei Li, W Berry Lyons, Patrick McGill, James McManis, David McPike, Anatoly Mironov, Molly Patterson, Graham Roberts, James Rot, Cathy Trainor, Martyn Tranter, John Winans, and Bob Zook. Biogeochemical and historical drivers of microbial community composition and structure in sediments from Mercer Subglacial Lake, West Antarctica. *ISME Communications*, 3(1), January 2023.

[6] Kyung Mo Kim, Kyuin Hwang, Hanbyul Lee, Ahnna Cho, Christina L. Davis, Brent C. Christner, John C. Priscu, and Ok-Sun Kim. Genetic isolation and metabolic complexity of an Antarctic subglacial microbiome. *Nature Communications*, 16(1), August 2025.

[7] Connor T Skennerton, Mohamed F Haroon, Ariane Briegel, Jian Shi, Grant J Jensen, Gene W Tyson, and Victoria J Orphan. Phylogenomic analysis of Candidatus 'Izimaplasma' species: free-living representatives from a Tenericutes clade found in methane seeps. *The ISME Journal*, 10(11):2679–2692, April 2016.

[8] Susannah J Salter, Michael J Cox, Elena M Turek, Szymon T Calus, William O Cookson, Miriam F Moffatt, Paul Turner, Julian Parkhill, Nicholas J Loman, and Alan W Walker. Reagent and laboratory contamination can critically impact sequence-based microbiome analyses. *BMC Biology*, 12(1), November 2014.

[9] J.A. Mikucki, C.G. Schuler, I. Digel, J. Kowalski, M.J. Tuttle, M. Chua, R. Davis, A.M. Purcell, D. Ghosh, G. Francke, M. Feldmann, C. Espe, D. Heinen, B. Dachwald, J. Clemens, W.B. Lyons, and S. Tulaczyk. Field-based planetary protection operations for melt probes: Validation of clean access into the Blood Falls, Antarctica, englacial ecosystem. *Astrobiology*, 23(11):1165–1178, November 2023.

[10] Donovan H Parks, Pierre-Alain Chaumeil, Aaron J Mussig, Christian Rinke, Maria Chuvochina, and Philip Hugenholtz. GTDB release 10: a complete and systematic taxonomy for 715230 bacterial and 17245 archaeal genomes. *Nucleic Acids Research*, October 2025.

[11] Tyler P. Barnum, Alexander Crits-Christoph, Michael Molla, Paul Carini, Henry H. Lee, and Nili Ostrov. Predicting microbial growth conditions from amino acid composition. *bioRXiv*, March 2024.

[12] Silvia Frisia, Laura S. Weyrich, John Hellstrom, Andrea Borsato, Nicholas R. Golledge, Alexandre M. Anesio, Petra Bajo, Russell N. Drysdale, Paul C. Augustinus, Camille Rivard, and Alan Cooper. The influence of Antarctic subglacial volcanism on the global iron cycle during the Last Glacial Maximum. *Nature Communications*, 8(1), June 2017.

[13] Kirsten A. Ziesemer, Allison E. Mann, Krithivasan Sankaranarayanan, Hannes Schroeder, Andrew T. Ozga, Bernd W. Brandt, Egija Zaura, Andrea Waters-Rist, Menno Hoogland, Domingo C. Salazar-García, Mark Aldenderfer, Camilla Speller, Jessica Hendy, Darlene A. Weston, Sandy J. MacDonald, Gavin H. Thomas, Matthew J. Collins, Cecil M. Lewis, Corinne Hofman, and Christina Warinner. Intrinsic challenges in ancient microbiome reconstruction using 16S rRNA gene amplification. *Scientific Reports*, 5(1), November 2015.

[14] Mario Fernández, Salvador Barahona, Fernando Gutierrez, Jennifer Alcaíno, Víctor Cifuentes, and Marcelo Baeza. Bacterial diversity, metabolic profiling, and application potential of Antarctic soil metagenomes. *Current Issues in Molecular Biology*, 46(11):13165–13178, November 2024.

[15] Claudia Coleine, Davide Albanese, Angelique E. Ray, Manuel Delgado-Baquerizo, Jason E. Stajich, Timothy J. Williams, Stefano Larsen, Susannah Tringe, Christa Pennacchio, Belinda C. Ferrari, Claudio Donati, and Laura Selbmann. Metagenomics untangles potential adaptations of Antarctic endolithic bacteria at the fringe of habitability. *Science of The Total Environment*, 917:170290, March 2024.

[16] Linnea F M Kop, Hanna Koch, Mike S M Jetten, Holger Daims, and Sebastian Lücker. Metabolic and phylogenetic diversity in the phylum nitrospinota revealed by comparative genome analyses. *ISME Communications*, 4(1), January 2024.

[17] Maliheh Mehrshad, Margarita Lopez-Fernandez, John Sundh, Emma Bell, Domenico Simone, Moritz Buck, Rizlan Bernier-Latmani, Stefan Bertilsson, and Mark Dopson. Energy efficiency and biological interactions define the core microbiome of deep oligotrophic groundwater. *Nature Communications*, 12(1), July 2021.

[18] Bianca De Sanctis, Cade Mirchandi, Haoran Dong, Ruairidh MacLeod, Russ Corbett-Detig, and Yucheng Wang. Bamdam: A post-mapping authentication toolkit for ancient metagenomics. *Genome Biology*, In Press. 2025. GitHub: https://github.com/bdesanctis/bamdam/.

[19] Timothy D'Angelo, Jacqueline Goordial, Melody R Lindsay, Julia McGonigle, Anne Booker, Duane Moser, Ramunas Stepanauskus, and Beth N Orcutt. Replicated life-history patterns and subsurface origins of the bacterial sister phyla nitrospirota and nitrospinota. *The ISME Journal*, 17(6):891–902, April 2023.

[20] T. Blackburn, G. H. Edwards, S. Tulaczyk, M. Scudder, G. Piccione, B. Hallet, N. McLean, J. C. Zachos, B. Cheney, and J. T. Babbe. Ice retreat in Wilkes Basin of East Antarctica during a warm interglacial. *Nature*, 583(7817):554–559, July 2020.

[21] Graham H. Edwards, Terrence Blackburn, Gavin Piccione, Slawek Tulaczyk, Gifford H. Miller, and Cosmo Sikes. Terrestrial evidence for ocean forcing of Heinrich events and subglacial hydrologic connectivity of the Laurentide Ice Sheet. *Science Advances*, 8(42), October 2022.

[22] Gavin Piccione, Terrence Blackburn, Slawek Tulaczyk, E. Troy Rasbury, Mathis P. Hain, Daniel E. Ibarra, Katharina Methner, Chloe Tinglof, Brandon Cheney, Paul Northrup, and Kathy Licht. Subglacial precipitates record Antarctic ice sheet response to late Pleistocene millennial climate cycles. *Nature Communications*, 13(1), September 2022.

[23] Gavin Piccione, Terrence Blackburn, Paul Northrup, Slawek Tulaczyk, and Troy Rasbury. Antarctic subglacial trace metal mobility linked to climate change across termination III. *The Cryosphere*, 19(6):2247–2261, June 2025.

[24] Jessica Gagliardi, Terrence Blackburn, Gavin Piccione, Slawek Tulaczyk, and C. Brenhin Keller. Subglacial precipitates record Antarctic Ice Sheet response to Southern Ocean warming. *Geophysical Research Letters*, 52(8), April 2025.

[25] L. Bazin, A. Landais, B. Lemieux-Dudon, H. Toyé Mahamadou Kele, D. Veres, F. Parrenin, P. Martinerie, C. Ritz, E. Capron, V. Lipenkov, M.-F. Loutre, D. Raynaud, B. Vinther, A. Svensson, S. O. Rasmussen, M. Severi, T. Blunier, M. Leuenberger, H. Fischer, V. Masson-Delmotte, J. Chappellaz, and E. Wolff. An optimized multi-proxy, multi-site Antarctic ice and gas orbital chronology (aicc2012): 120–800 ka. *Climate of the Past*, 9(4):1715–1731, August 2013.

[26] Wei Shen, Botond Sipos, and Liuyang Zhao. SeqKit2: A Swiss army knife for sequence and alignment processing. *iMeta*, 3(3), April 2024.

[27] Yu Peng, Henry C. M. Leung, S. M. Yiu, and Francis Y. L. Chin. IDBA-UD: a de novo assembler for single-cell and metagenomic sequencing data with highly uneven depth. *Bioinformatics*, 28(11):1420–1428, April 2012.

[28] Doug Hyatt, Gwo-Liang Chen, Philip F LoCascio, Miriam L Land, Frank W Larimer, and Loren J Hauser. Prodigal: prokaryotic gene recognition and translation initiation site identification. *BMC Bioinformatics*, 11(1), March 2010.

[29] Carlos P Cantalapiedra, Ana Hernández-Plaza, Ivica Letunic, Peer Bork, and Jaime Huerta-Cepas. eggnog-mapper v2: Functional annotation, orthology assignments, and domain prediction at the metagenomic scale. *Molecular Biology and Evolution*, 38(12):5825–5829, October 2021.

[30] Benjamin Buchfink, Chao Xie, and Daniel H Huson. Fast and sensitive protein alignment using DIAMOND. *Nature Methods*, 12(1):59–60, November 2014.

[31] Heng Li and Richard Durbin. Fast and accurate short read alignment with burrows–wheeler transform. *Bioinformatics*, 25(14):1754–1760, May 2009.

[32] Adrien Oliva, Raymond Tobler, Bastien Llamas, and Yassine Souilmi. Additional evaluations show that specific bwa-aln settings still outperform bwa-mem for ancient dna data alignment. *Ecology and Evolution*, 11(24):18743–18748, December 2021.

[33] Nicholas B. Dragone, Jessica B. Henley, Hannah Holland-Moritz, Melisa Diaz, Ian D. Hogg, W. Berry Lyons, Diana H. Wall, Byron J. Adams, and Noah Fierer. Elevational constraints on the composition and genomic attributes of microbial communities in Antarctic soils. *mSystems*, 7(1), February 2022.

[34] Sean K. Bay, Xiyang Dong, James A. Bradley, Pok Man Leung, Rhys Grinter, Thanavit Jirapanjawat, Stefan K. Arndt, Perran L. M. Cook, Douglas E. LaRowe, Philipp A. Nauer, Eleonora Chiri, and Chris Greening. Trace gas oxidizers are widespread and active members of soil microbial communities. *Nature Microbiology*, 6(2):246–256, January 2021.

[35] Xiaoli Yu, Jiayin Zhou, Wen Song, Mengzhao Xu, Qiang He, Yisheng Peng, Yun Tian, Cheng Wang, Longfei Shu, Shanquan Wang, Qingyun Yan, Jihua Liu, Qichao Tu, and Zhili He. Scycdb: A curated functional gene database for metagenomic profiling of sulphur cycling pathways. *Molecular Ecology Resources*, 21(3):924–940, December 2020.

[36] Qichao Tu, Lu Lin, Lei Cheng, Ye Deng, and Zhili He. Ncycdb: a curated integrative database for fast and accurate metagenomic profiling of nitrogen cycling genes. *Bioinformatics*, 35(6):1040–1048, August 2018.

[37] Lu Qian, Xiaoli Yu, Jiayin Zhou, Hang Gu, Jijuan Ding, Yisheng Peng, Qiang He, Yun Tian, Jihua Liu, Shanquan Wang, Cheng Wang, Longfei Shu, Qingyun Yan, Jianguo He, Guangli Liu, Qichao Tu, and Zhili He. MCycDB: A curated database for comprehensively profiling methane cycling processes of environmental microbiomes. *Molecular Ecology Resources*, 22(5):1803–1823, February 2022.

[38] Arkadiy I. Garber, Kenneth H. Nealson, Akihiro Okamoto, Sean M. McAllister, Clara S. Chan, Roman A. Barco, and Nancy Merino. Fegenie: A comprehensive tool for the identification of iron genes and iron gene neighborhoods in genome and metagenome assemblies. *Frontiers in Microbiology*, 11, January 2020.

[39] Arren Bar-Even, Elad Noor, and Ron Milo. A survey of carbon fixation pathways through a quantitative lens. *Journal of Experimental Botany*, 63(6):2325–2342, December 2011.

[40] The UniProt Consortium. Uniprot: a worldwide hub of protein knowledge. *Nucleic Acids Research*, 47(D1):D506–D515, November 2018.

[41] J. A. Mikucki, P. A. Lee, D. Ghosh, A. M. Purcell, A. C. Mitchell, K. D. Mankoff, A. T. Fisher, S. Tulaczyk, S. Carter, M. R. Siegfried, H. A. Fricker, T. Hodson, J. Coenen, R. Powell, R. Scherer, T. Vick-Majors, A. A. Achberger, B. C. Christner, and M. Tranter. Subglacial Lake Whillans microbial biogeochemistry: a synthesis of current knowledge. *Philosophical Transactions of the Royal Society A: Mathematical, Physical and Engineering Sciences*, 374(2059):20140290, January 2016.

[42] Nicholas B. Dragone, Kerry Whittaker, Olivia M. Lord, Emily A. Burke, Helen Dufel, Emily Hite, Farley Miller, Gabrielle Page, Dan Slayback, and Noah Fierer. The early microbial colonizers of a short-lived volcanic island in the kingdom of tonga. *mBio*, 14(1), February 2023.

[43] Maximiliano Ortiz, Pok Man Leung, Guy Shelley, Thanavit Jirapanjawat, Philipp A. Nauer, Marc W. Van Goethem, Sean K. Bay, Zahra F. Islam, Karen Jordaan, Surendra Vikram, Steven L. Chown, Ian D. Hogg, Thulani P. Makhalanyane, Rhys Grinter, Don A. Cowan, and Chris Greening. Multiple energy sources and metabolic strategies sustain microbial diversity in Antarctic desert soils. *Proceedings of the National Academy of Sciences*, 118(45), November 2021.

[44] Sergey Nurk, Dmitry Meleshko, Anton Korobeynikov, and Pavel A. Pevzner. metaSPAdes: a new versatile metagenomic assembler. *Genome Research*, 27(5):824–834, March 2017.

[45] Louis Kraft, Johannes Söding, Martin Steinegger, Annika Jochheim, Peter Wad Sackett, Antonio Fernandez-Guerra, and Gabriel Renaud. Carpedeam: Ade novometagenome assembler for heavily damaged ancient datasets. August 2024.

[46] Ben Langmead and Steven L Salzberg. Fast gapped-read alignment with bowtie 2. *Nature Methods*, 9(4):357–359, March 2012.

[47] Petr Danecek, James K Bonfield, Jennifer Liddle, John Marshall, Valeriu Ohan, Martin O Pollard, Andrew Whitwham, Thomas Keane, Shane A McCarthy, Robert M Davies, and Heng Li. Twelve years of SAMtools and BCFtools. *GigaScience*, 10(2), 02 2021. giab008.

[48] A. Murat Eren, Evan Kiefl, Alon Shaiber, Iva Veseli, Samuel E. Miller, Matthew S. Schechter, Isaac Fink, Jessica N. Pan, Mahmoud Yousef, Emily C. Fogarty, Florian Trigodet, Andrea R. Watson, Özcan C. Esen, Ryan M. Moore, Quentin Clayssen, Michael D. Lee, Veronika Kivenson, Elaina D. Graham, Bryan D. Merrill, Antti Karkman, Daniel Blankenberg, John M. Eppley, Andreas Sjödin, Jarrod J. Scott, Xabier Vázquez-Campos, Luke J. McKay, Elizabeth A. McDaniel, Sarah L. R. Stevens, Rika E. Anderson, Jessika Fuessel, Antonio Fernandez-Guerra, Lois Maignien, Tom O. Delmont, and Amy D. Willis. Community-led, integrated, reproducible multi-omics with anvi'o. *Nature Microbiology*, 6(1):3–6, December 2020.

[49] Johannes Alneberg, Brynjar Smári Bjarnason, Ino de Bruijn, Melanie Schirmer, Joshua Quick, Umer Z Ijaz, Leo Lahti, Nicholas J Loman, Anders F Andersson, and Christopher Quince. Binning metagenomic contigs by coverage and composition. *Nature Methods*, 11(11):1144–1146, September 2014.

[50] Yu-Wei Wu, Blake A. Simmons, and Steven W. Singer. Maxbin 2.0: an automated binning algorithm to recover genomes from multiple metagenomic datasets. *Bioinformatics*, 32(4):605–607, October 2015.

[51] Dongwan D. Kang, Feng Li, Edward Kirton, Ashleigh Thomas, Rob Egan, Hong An, and Zhong Wang. Metabat 2: an adaptive binning algorithm for robust and efficient genome reconstruction from metagenome assemblies. *PeerJ*, 7:e7359, July 2019.

[52] Christian M. K. Sieber, Alexander J. Probst, Allison Sharrar, Brian C. Thomas, Matthias Hess, Susannah G. Tringe, and Jillian F. Banfield. Recovery of genomes from metagenomes via a dereplication, aggregation and scoring strategy. *Nature Microbiology*, 3(7):836–843, May 2018.

[53] Donovan H. Parks, Michael Imelfort, Connor T. Skennerton, Philip Hugenholtz, and Gene W. Tyson. Checkm: assessing the quality of microbial genomes recovered from isolates, single cells, and metagenomes. *Genome Research*, 25(7):1043–1055, May 2015.

[54] Pierre-Alain Chaumeil, Aaron J Mussig, Philip Hugenholtz, and Donovan H Parks. GTDB-Tk v2: memory friendly classification with the genome taxonomy database. *Bioinformatics*, 38(23):5315–5316, October 2022.

[55] Donovan H Parks, Maria Chuvochina, Christian Rinke, Aaron J Mussig, Pierre-Alain Chaumeil, and Philip Hugenholtz. Gtdb: an ongoing census of bacterial and archaeal diversity through a phylogenetically consistent, rank normalized and complete genome-based taxonomy. *Nucleic Acids Research*, 50(D1):D785–D794, September 2021.

[56] Michael Shaffer, Mikayla A Borton, Bridget B McGivern, Ahmed A Zayed, Sabina Leanti La Rosa, Lindsey M Solden, Pengfei Liu, Adrienne B Narrowe, Josué Rodríguez-Ramos, Benjamin Bolduc, M Consuelo Gazitúa, Rebecca A Daly, Garrett J Smith, Dean R Vik, Phil B Pope, Matthew B Sullivan, Simon Roux, and Kelly C Wrighton. Dram for distilling microbial metabolism to automate the curation of microbiome function. *Nucleic Acids Research*, 48(16):8883–8900, August 2020.
